# Micro-C XL: assaying chromosome conformation at length scales from the nucleosome to the entire genome

**DOI:** 10.1101/071357

**Authors:** Tsung-Han S. Hsieh, Geoffrey Fudenberg, Anton Goloborodko, Oliver J. Rando

**Affiliations:** Department of Biochemistry and Molecular Pharmacology, University of Massachusetts Medical School, Worcester, MA 01605, USA; Department of Physics and Center for 3D Structure and Physics of the Genome, Massachusetts Institute of Technology (MIT), Cambridge, MA 02139, USA

## Abstract

Structural analysis of chromosome folding in vivo has been revolutionized by Chromosome Conformation Capture (3C) and related methods, which use proximity ligation to identify chromosomal loci in physical contact. We recently described a variant 3C technique, Micro-C, in which chromatin is fragmented to mononucleosomes using micrococcal nuclease, enabling nucleosome-resolution folding maps of the genome. Here, we describe an improved Micro-C protocol using long crosslinkers, termed Micro-C XL, which exhibits greatly increased signal to noise, and provides further insight into the folding of the yeast genome. We also find that signal to noise is much improved in Micro-C XL libraries generated from relatively insoluble chromatin as opposed to soluble material, providing a simple method to physically enrich for bona-fide long-range interactions. Micro-C XL maps of the budding and fission yeast genomes reveal both short-range chromosome fiber features such as chromosomally-interacting domains (CIDs), as well as higher-order features such as clustering of centromeres and telomeres, thereby addressing the primary discrepancy between prior Micro-C data and reported 3C and Hi-C analyses. Interestingly, comparison of chromosome folding maps of *S. cerevisiae* and *S. pombe* revealed widespread qualitative similarities, yet quantitative differences, between these distantly-related species. Micro-C XL thus provides a single assay suitable for interrogation of chromosome folding at length scales from the nucleosome to the full genome.

## INTRODUCTION

The compaction and organization of the physical genome has wide-ranging consequences for genomic function^1-4^. In eukaryotes, the first level of genome compaction is organization into the characteristic “beads on a string” structure, with nucleosomes separated by relatively accessible linker DNA. Our understanding of this primary structure of chromatin is well-developed, with multiple crystal structures solved for the nucleosome^5, 6^, and a plethora of genome-wide studies that identify the positions of individual nucleosomes across the genome in various organisms, in some cases at single nucleotide-resolution^7^. The next step in chromosome folding remains relatively poorly-characterized; for example, the long-held belief that chromatin fibers form a helical secondary structure termed the 30 nm fiber is increasingly subject to debate^8-15^.

Structural analysis of chromosome folding beyond the nucleosome fiber has been revolutionized by the Chromosome Conformation Capture (3C) family of techniques^2, 16^. In 3C-based protocols, chromatin is first crosslinked in vivo using formaldehyde to capture physical interactions between distal regions of the genome. Chromatin is subsequently fragmented, and ligation of chromatin fragments is used to generate chimeric DNA molecules. Sequencing these molecular libraries provides a readout of genomic loci that were crosslinked to one another via protein-protein interactions. Genome-wide variants of 3C, such as Hi-C, have revealed a number of organizational features of eukaryotic genomes at increasingly fine resolutions, from the scale of full chromosomal territories, to multi-Mb active and inactive compartments, to hundred-kb contact domains (TADs), to enhancer-promoter loops^17-28^. While many factors impact the effective resolution of a 3C/Hi-C dataset, including sequencing depth and library complexity^29^, a fundamental limit to genomic resolution is the size of the fragments generated before physical interactions are captured via ligation. Since the majority of 3C-based experiments rely on restriction enzymes for fragmentation of the genome – resulting in genomic fragments that are both long relative to the nucleosome, and inhomogeneously spaced along the genome – current Hi-C datasets are limited to ∼1 kb resolution.

To improve the resolution of 3C-based techniques, we recently developed a high resolution 3C-based technique, dubbed “Micro-C”, in which fragmentation of the genome is accomplished using micrococcal nuclease (MNase) to enable mononucleosome-resolution analysis of chromosome folding^17^. While the improved resolution afforded by Micro-C enabled the identification of features such as chromosomally-interacting domains – “CIDs” – in budding yeast that had not previously been discernible using a restriction enzyme-based 3C technique^30^, known higher-order interactions such as centromere clustering were poorly recovered using our technique. In addition, although several studies have reported “gene loops” in budding yeast using 3C methods^31, 32^, we found no evidence for gene loops using Micro-C.

These discrepancies with the literature motivated a deeper exploration of the effects of specific protocol steps on the results of Micro-C analysis of chromosome folding. Most notably, we sought to determine whether the reliance on formaldehyde, a “zero length” crosslinker, to crosslink genomic loci to one another might limit the ability of 3C-related methods to fully interrogate chromosome structure. To investigate whether longer crosslinkers might reveal additional features of local chromatin structure, we characterized the effects of two long protein-protein crosslinkers on Micro-C maps of the budding yeast genome. A revised Micro-C protocol incorporating long crosslinkers, which we named “Micro-C XL”, not only recapitulated the local chromatin structures previously revealed by Micro-C, but also robustly recovered higher-order features such as centromere-centromere interactions. Micro-C XL thus overcomes the key technical limitation of the original Micro-C protocol, providing a single protocol for analysis of chromosomal folding from the scale of nucleosomes to the full genome. We also characterized Micro-C XL profiles in pellet and supernatant fractions of crosslinked chromatin, finding that chromatin contacts are enriched in relatively insoluble chromatin, thereby providing a simple technical approach to improve signal-to-noise in Micro-C maps. Finally, we compared Micro-C XL maps from *S. cerevisiae* and *S. pombe*, finding a general conservation of gene-scale folding behavior in these distantly-related species. Taken together, our results provide an updated Micro-C protocol for characterization of chromosome folding at all length scales, and provide additional high resolution insights into chromosome structure in two key model organisms.

## RESULTS

### Optimization of crosslinking conditions for Micro-C

We recently detailed a modified Hi-C^19^ protocol, termed Micro-C, in which micrococcal nuclease (MNase) digestion of crosslinked chromatin enables the analysis of chromosome folding at mononucleosomal resolution^17^. Our reported Micro-C maps robustly captured short-range interactions such as chromosomally-interacting domains (CIDs) in budding yeast, but exhibited poor recovery of higher-order features such as the centromere-centromere (CEN-CEN) and telomere-telomere (TEL-TEL) interactions that are well-known features of yeast genome organization. We reasoned that the use of the short-range crosslinker formaldehyde in 3C protocols might limit the chromosomal interactions captured due to the extreme physical proximity required for formaldehyde to crosslink two proteins.

We therefore sought to identify alternative crosslinkers or crosslinking conditions to improve Micro-C capture of CEN-CEN and other long-range interactions. Using qPCR primers designed to assay interactions either within the contact domain associated with *MDJ1* or between pairs of centromeres (**Fig. S1**), we tested a variety of different protein-protein crosslinkers and crosslinking conditions to identify conditions that best enabled recovery of longer-range (greater than ∼1 kb) interactions. These analyses identified two protein-protein crosslinkers that appeared to more efficiently crosslink distant nucleosomes within the *MDJ1* CID and to more efficiently capture CEN-CEN interactions – disuccinimidyl glutarate (DSG, a 7.7Å crosslinker), and ethylene glycol bis(succinimidyl succinate) (EGS, a 16.1Å crosslinker) (**Fig. 1a-b**). The improvements in signal-to-noise afforded by DSG and EGS were not observed when DSG or EGS were added prior to cell permabilization (not shown), consistent with our expectation that these molecules are too large to cross the yeast cell wall ^33^. In addition, improved signal-to-noise was not observed following crosslinking with higher concentrations of formaldehyde, longer incubation times, or when a second round of formaldehyde crosslinking was carried out after cell wall digestion (**Fig. S1e-f**), demonstrating that the improvements in the Micro-C protocol required some specific aspect of the DSG and EGS crosslinkers, rather than, say, an increase in the sheer density of crosslinks introduced into chromatin.

**Fig. 1.**
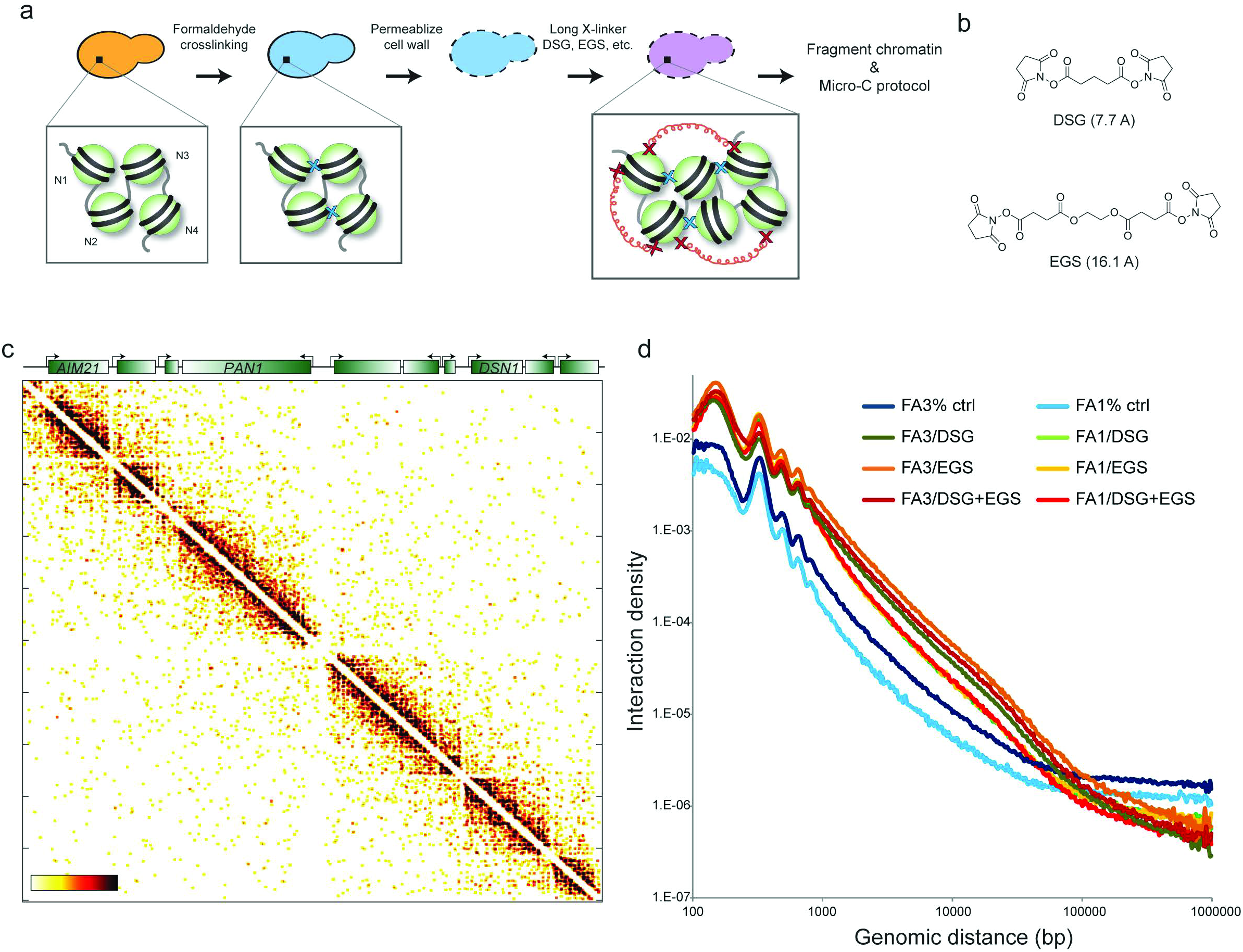
Overview of Micro-C XL. (**a**) Outline of changes to the Micro-C protocol. After budding yeast are fixed with formaldehyde, cells are permeabilized, then treated with one of several additional protein-protein crosslinkers. Crosslinked chromatin is then digested to mononucleosomes using micrococcal nuclease. End digestion and repair is used to introduce biotinylated nucleotides into mononucleosomal ends, and nucleosomes crosslinked to one another are ligated together at high dilution. Ligation products are then purified via streptavidin capture and size selection of dinucleosomesized DNA, and paired-end deep sequencing is used to characterize internucleosomal interactions genome-wide. (**b**) Structures of the two protein-protein crosslinkers used in Micro-C XL. (**c**) Example of Micro-C XL contact map for a 20 kb genomic stretch. The Micro-C XL protocol effectively recovers the chromosomally-interacting domains previously observed in Micro-C data^17^. (**d**) Plot of interaction density for all unidirectional (“IN-OUT”) read pairs, expressed as a fraction of potential pairwise interactions (per bp^2^) (y axis, log10), vs. genomic distance (x axis, log10) for various Micro-C protocols scaled to 10^9^ interactions.

We incorporated each of these longer crosslinkers into an altered Micro-C protocol, which we dubbed Micro-C XL (<u>MICRO</u>coccal nuclease-based analysis of <u>C</u>hromosome folding using long <u>X-L</u>inkers), and then sought to identify those features of yeast chromosome folding uniquely revealed using these crosslinkers (**Methods**). Briefly, actively growing budding yeast cultures are crosslinked with formaldehyde alone, formaldehyde + DSG, formaldehyde + EGS, or all three crosslinkers, and resulting chromatin is fragmented to mononucleosomes using MNase digestion (**Fig. 1a**). Crosslinked chromatin is then treated with T4 DNA polymerase in the absence of dNTPs to promote exonuclease activity. This leaves single stranded DNA ends, which are then repaired and biotinylated upon the addition of dNTPs, including biotin-dATP and biotin-dCTP, to the T4 polymerase reaction. Following DNA ligation, ligation products are purified away from unligated mononucleosomal DNA based 1) on ligation-dependent protection of biotinylated DNA from exonuclease attack, and 2) on size selection specifically of dinucleosome-sized ligation products. In practice, nucleosomal ligation products are first treated with exonuclease III to remove biotinylated nucleotides from free DNA ends, leaving biotinylated nucleotides specifically in nucleosomal ends that had been ligated to one another and thereby protected from exonuclease attack. DNA is then purified from deproteinated chromatin, and dinucleosome-sized ligation products are gel-purified away from unligated mononucleosomal DNA. Recovered DNA is then further purified on streptavidin beads to isolate only DNA carrying biotinylated nucleotides at ligation junctions that had been protected from exonuclease digestion. Purified ligation products are then used to generate deep sequencing libraries, and subject to Illumina paired-end sequencing using standard methods.

### Genome-wide analysis of chromosome folding by Micro-C XL

To investigate whether adding DSG or EGS to the Micro-C protocol provided additional insights into chromosome folding, we generated genome-wide Micro-C maps for budding yeast subject to a variety of crosslinking conditions. These conditions include 1% or 3% formaldehyde (FA) alone, or FA (1% or 3%) plus DSG, EGS, or both crosslinkers. Also, as described above, we generated similar datasets in which DSG or EGS were added prior to cell wall digestion as a negative control, as these molecules are not expected to cross the cell wall in budding yeast. Below, we primarily focus on the results using 3% FA with or without DSG and EGS, but results of other crosslinking conditions are noted when relevant.

In general, all four conditions (FA, FA/DSG, FS/EGS, and FA/DSG/EGS) yielded qualitatively similar results at the scale of individual genes, with chromosomal interaction domains of varying strength covering ∼1-5 genes (**Fig. 1c, S2**). Although CIDs were clearly observed in all four conditions, the addition of longer crosslinkers to the Micro-C protocol resulted in improved ability to visualize these structures (**Fig. S2** and see below). Importantly, as previously observed with Micro-C, we again found no evidence for a regular organization of the chromatin fiber above the nucleosomal scale, which would have manifested as a peak in interaction density at a genomic distance corresponding to the fiber size (**Fig. 1d, Table S1**). Moreover, compared to standard Micro-C we found that Micro-C XL exhibited substantially higher signal-to-noise (**Fig. 1d, S3**), consistent with the q-PCR results in **Fig. S1**.

Beyond recapitulating the key aspects of chromosome folding previously revealed by Micro-C, Micro-C XL resolved additional details that were not apparent in prior Micro-C maps. Most interestingly, in contrast to standard Micro-C crosslinking conditions, all three long crosslinking conditions captured very robust CEN-CEN and TEL-TEL interactions characteristic of the Rabl configuration for interphase chromosomes (**Figs. 2, S4**)^24, 27, 30^. This finding thus resolves the primary qualitative discrepancy between prior Micro-C data and known features of genomic folding in yeast, while preserving the ability of Micro-C to interrogate chromatin interactions at the 2-10 nucleosome scale.

**Fig. 2.**
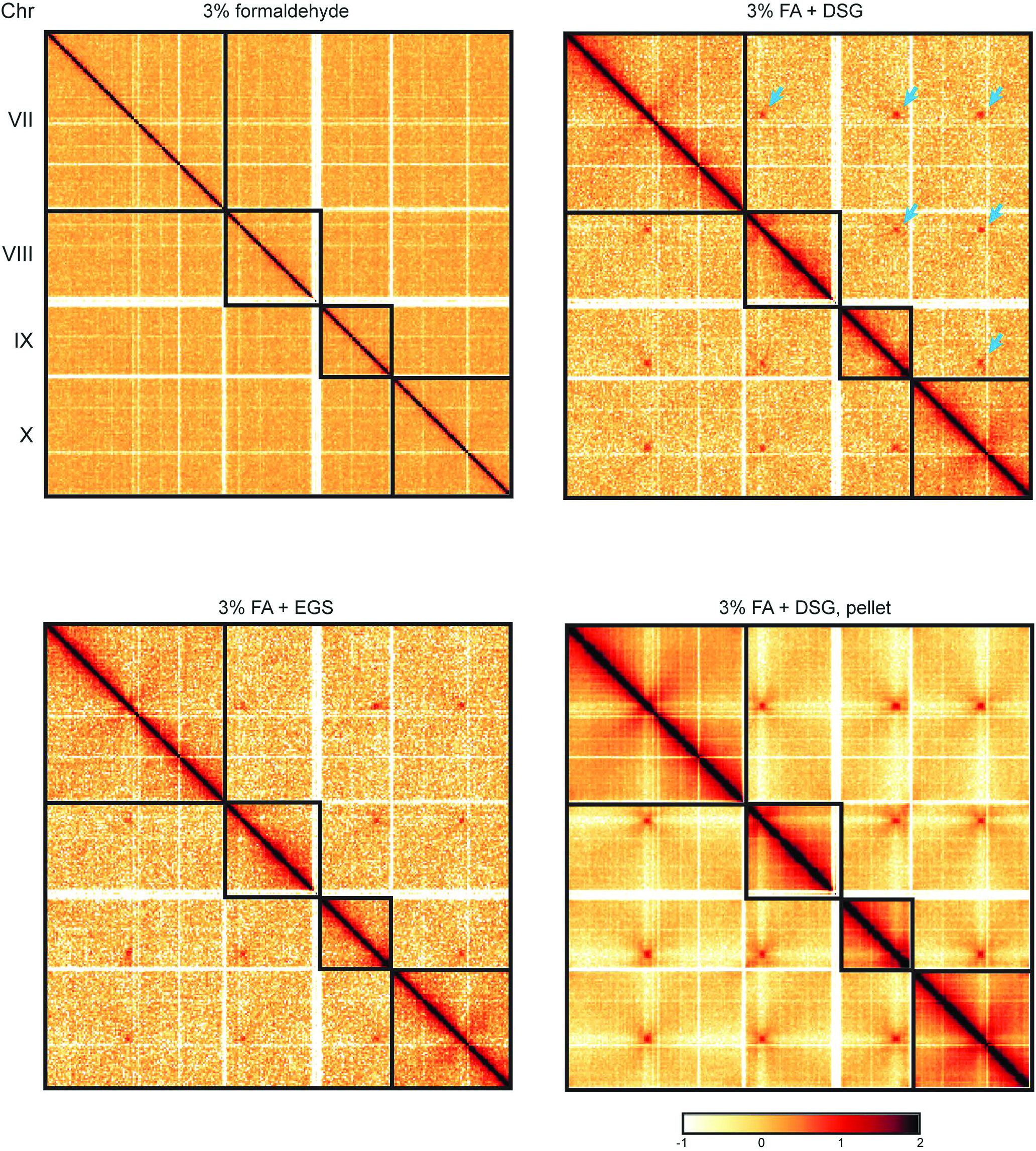
Micro-C XL robustly captures known interchromosomal interactions. Interaction maps for Micro-C data generated using 3% formaldehyde, 3% formaldehyde+DSG, or 3% formaldehyde+EGS are shown as indicated for budding yeast chromosomes VII through X. Here, data are normalized by read depth only and displayed as log10(counts per million). Lower right panel shows Micro-C XL data for 3% formaldehyde+DSG in which crosslinked chromatin was subject to centrifugation following MNase digestion, and the pellet fraction was subject to downstream protocol steps (see also **Fig. 3**). For 3% FA+DSG, CEN-CEN interactions are shown with blue arrows. As previously observed, telomere-telomere interactions are only observed between a subset of chromosome arms (which do not include interactions between chromosomes VII to X) in budding yeast.

**Fig. 3.**
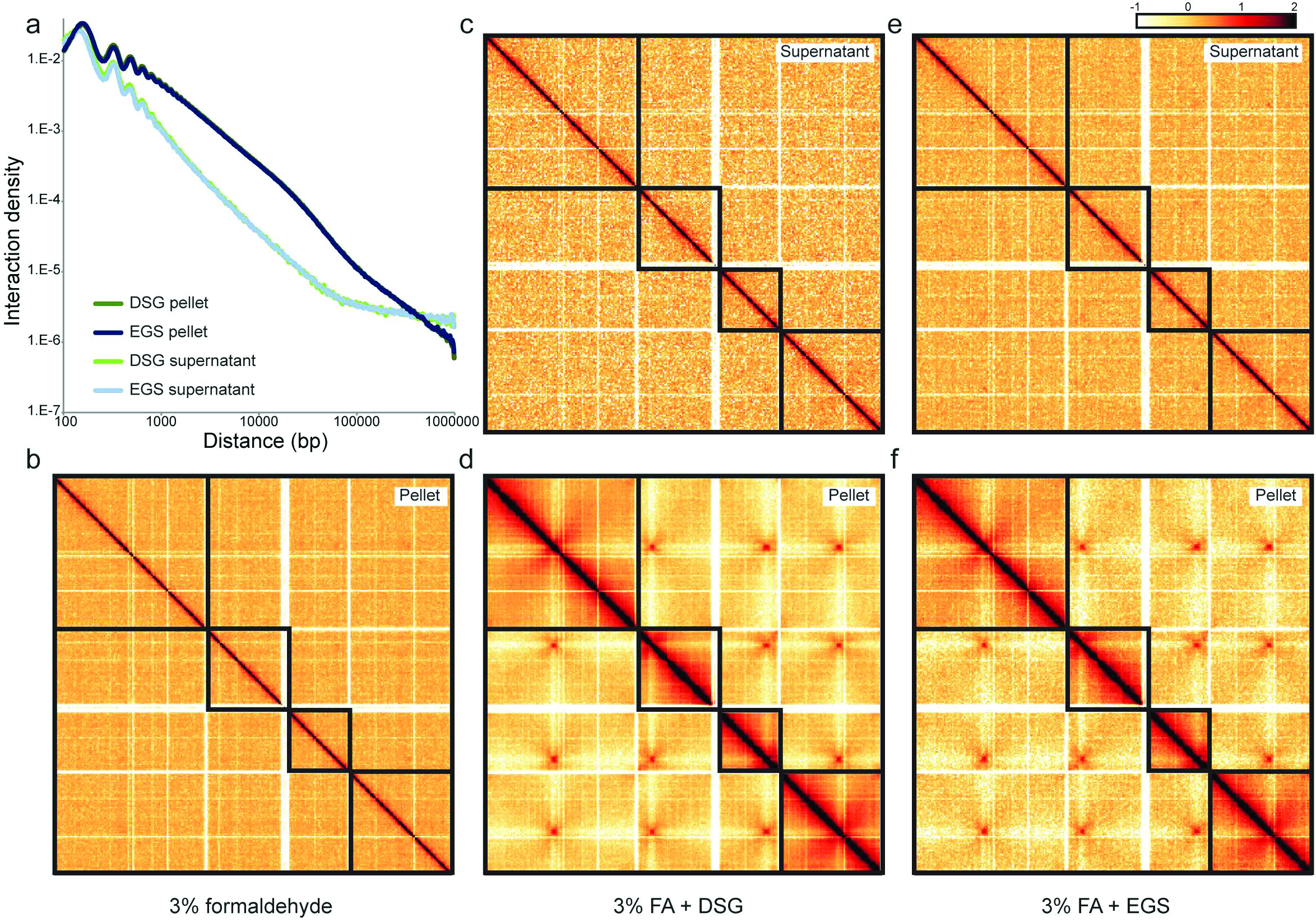
Micro-C XL interactions are enriched in insoluble chromatin. (**a**) Plot of interaction density (y axis, log10) vs. genomic distance (x axis, log10) for four Micro-C XL libraries, normalized as in **Fig. 1d**. One pair of libraries were crosslinked with 3% FA+DSG, then MNase-digested chromatin was separated into soluble and insoluble fractions by centrifugation; the same procedure was also repeated for yeast crosslinked with 3% FA+EGS. In both cases, the relatively soluble Micro-C library exhibited far lower signal to noise, with relatively rapid decay of interactions with increasing distance, compared to libraries constructed from pellet material. (**b-f**) Micro-C interaction maps for chromosomes VII to X are shown as in **Fig. 2**, for pellet (**b**, **d**, **f**) and supernatant (**c**, **e**) libraries as indicated. Note that the strong enrichment of CEN-CEN interactions in the pellet fraction requires long-distance crosslinkers, as it is not observed for 3% FA chromatin pellets (**b**).

We conclude that both DSG and EGS dramatically extend the length scale at which chromosome folding can be assayed by Micro-C, enabling analysis at scales from the local chromatin fiber to the full genome. Interestingly, this improvement can largely be ascribed to a decrease in the background levels of ligation between distant genomic regions (**Fig. S3, Table S1**) – the decrease in this “noise floor” seen using the Micro-C XL protocol is likely to account for the improved ability to measure relatively low-abundance signals such as CEN-CEN interactions.

### Bona-fide Micro-C contacts are primarily found in relatively insoluble chromatin

We next sought to uncover whether Micro-C data are affected by fractionation of crosslinked chromatin prior to proximity ligation. This was motivated by the absence of “gene loops”^31, 32^ in our previous Micro-C analysis – the 3C method used in several studies of gene loops includes a step in which insoluble chromatin is pelleted and isolated prior to ligation, and it is known that different chromatin structures are likely to be present in soluble vs. insoluble crosslinked chromatin^34^. We therefore carried out Micro-C XL in which fragmented chromatin was centrifuged after the completion of MNase digestion to separate soluble from insoluble chromatin (**Methods**), and proximity ligation was carried out separately on pellet and supernatant material (**Fig. 3**).

Micro-C XL maps from supernatant material were extremely noisy at longer distances, and did not identify known aspects of higher-order organization. In contrast, contact maps generated from relatively insoluble chromatin had excellent signal to noise, and robustly captured CEN-CEN interactions. We conclude that this reflects either preferential precipitation of crosslinked fragments, higher ligation efficiency in the pellet, or higher specificity of ligation in the pellet; i.e. noise in the supernatant dataset is elevated due to ligations in solution between freely-moving, likely uncrosslinked, nucleosomes causing artefactual contacts in trans. These related hypotheses are of course not mutually exclusive, and may all contribute to the improved signal to noise seen in Micro-C XL maps from pellet material (**Fig. S3**, right panel).

We next searched for evidence of gene loops^31, 32^ in the dataset generated from relatively insoluble chromatin. Here, we consider a gene “loop” to be characterized by an increased contact frequency between the gene start and stop relative to other locus pairs in their vicinity (similar to previous definitions of peaks in Hi-C contact maps –^28^), rather than a gene-wide increase in relative contact frequency. In general, it is clear that, for any given nucleosome, raw interaction counts (either normalized only for library depth, or normalized additionally for nucleosome occupancy) decay steadily with increasing distance and do not exhibit an uptick at gene ends, which is the signature of a looping interaction^35, 36^ – see example in **Fig. 1c**, or averaged “metagenes” in **Figs. S5-6**. Nevertheless, with our population-average contact maps we cannot rule out a scenario where populations of various length gene loops are formed dynamically over gene bodies (as proposed for enhancer-promoter interactions –^37^).

Although our data thus do not support the concept of widespread end-to-end gene loops at transcribed genes, visual inspection of Micro-C XL data did reveal a small number of possible looping interactions at multi-gene scale that were apparent even in interaction counts not normalized for distance (**Fig. S7**). Validation and functional analysis of these apparent loops will be the subject of future studies.

### Comparison of chromosome folding in *S. cerevisiae* and *S. pombe*

Although many aspects of chromosome folding are conserved between *S. cerevisiae* and other eukaryotes, *S. cerevisiae* lack several evolutionarily widespread chromatin regulatory systems, such as the H3K9me3/HP1 and H3K27me3/Polycomb systems for gene repression found in many eukaryotes. We therefore carried out Micro-C XL in the fission yeast *S. pombe* to ascertain the similarities and differences in chromosome folding between these distantly-related microbes, and to demonstrate the broad applicability of our methods. Key aspects of the Micro-C XL protocol proved equally important in fission yeast, as for example maps generated from relatively insoluble chromatin exhibited far less noise compared to maps based on soluble chromatin (**Fig. S8**). Overall, our data were well-correlated (Spearman’s r = 0.77 using 10 kb bins) with a prior Hi-C analysis of *S. pombe* chromatin by Mizuguchi *et al* (**Fig. S9**)^24^. As in budding yeast, Micro-C XL maps in *S. pombe* revealed frequent interactions along the diagonal, robust CEN-CEN and TEL-TEL interactions, and a depletion of interactions between centromeres and chromosome arms (**Fig. 4a**). We did find quantitative differences in such large-scale aspects of chromosome folding, as *S. pombe* chromosomes exhibited slightly stronger centromere clustering, and substantially stronger telomere clustering (**Fig. S10**). These differences do not appear to be a consequence of the profound cell cycle differences between budding and fission yeast (**Fig. S10c-d**), but could potentially be explained by any number of other features ranging from the smaller number of longer chromosomes in *S. pombe*, to the molecular details of interactions between pairs of H3K9-methylated nucleosomes present in this species but not in *S. cerevisiae*.

**Fig. 4.**
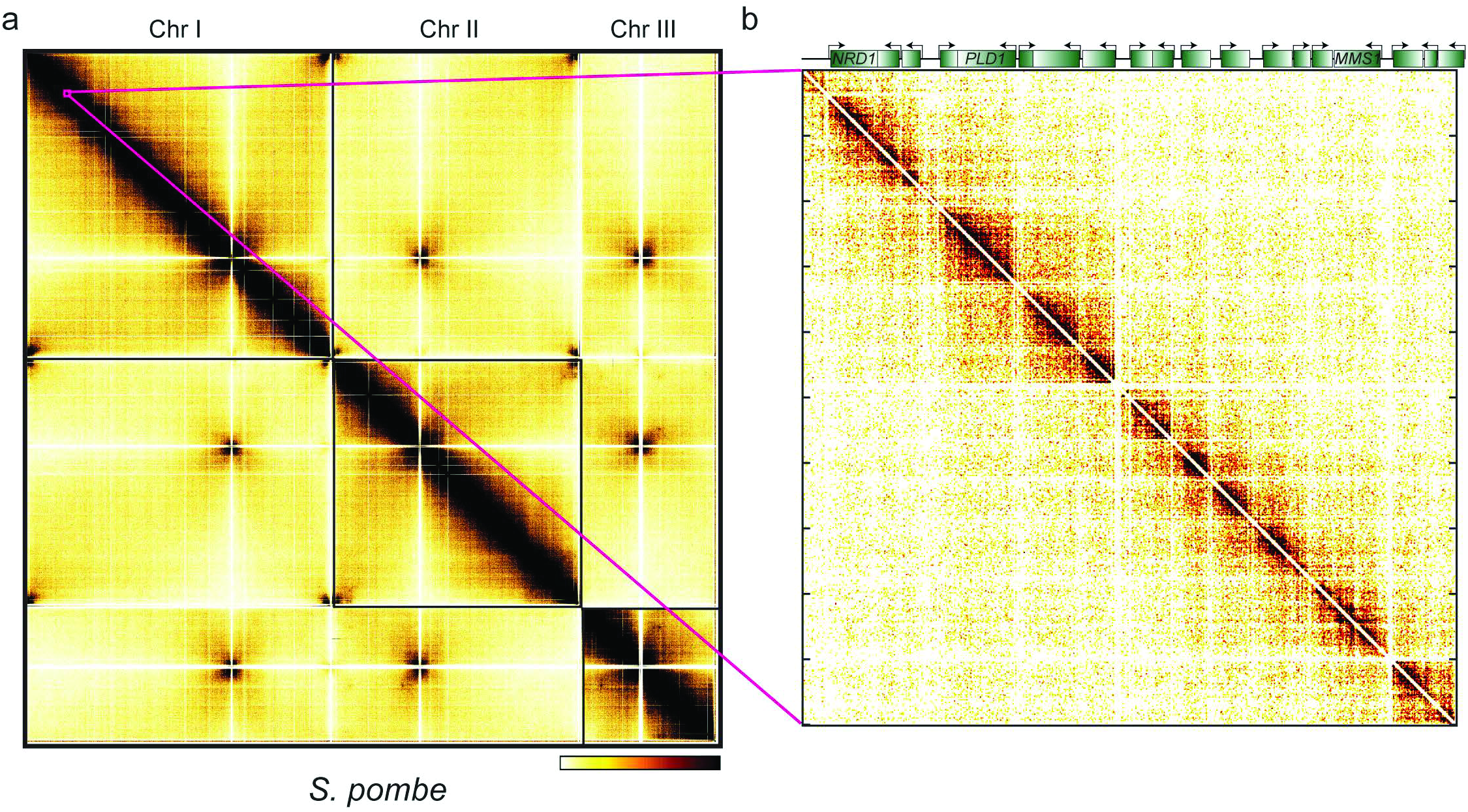
Analysis of chromosome folding in *S. pombe*. (**a**) Whole genome Micro-C XL interaction map for *S. pombe*. Here, data are shown for yeast crosslinked with 3% FA+DSG and pelleted prior to ligation. Similar results were obtained with 3% FA+EGS (not shown). Key features of this map include robust clustering of centromeres and telomeres (with the exception of the rDNA-carrying chromosome III telomeres, which were excluded from analysis based on their repetitive nature), and strong depletion of interactions between centromeres and chromosome arms. (**b**) Zoom-in on *S. pombe* Chr I:553,200-609,400, showing widespread contact domains typically associated with individual genes, but occasionally associated with blocks of ∼2-5 genes.

As prior studies of chromosome folding in *S. pombe* were performed with ∼10 kb kb resolution, we next turned to those aspects of chromatin structure uniquely interrogated using the enhanced resolution of Micro-C. Visual inspection of chromosome folding revealed abundant contact domains associated with ∼1-5 genes and separated by promoter regions (**Fig. 4b**), analogous to the CID structures in budding yeast. As in budding yeast, promoters in fission yeast acted as efficient boundaries between CIDs, and metagene analysis revealed remarkably similar behavior in both yeast species at the length scale of individual genes or promoters (**Fig. 5, S11-12**). In addition, plots of interaction frequency vs. distance, and distributions of contact domain length, were qualitatively similar, yet quantitatively different, in these two species; in particular, differences at short distances in the positions of interaction maxima correspond to known differences in nucleosome repeat length in these two species (**Fig. S13**).

**Fig. 5.**
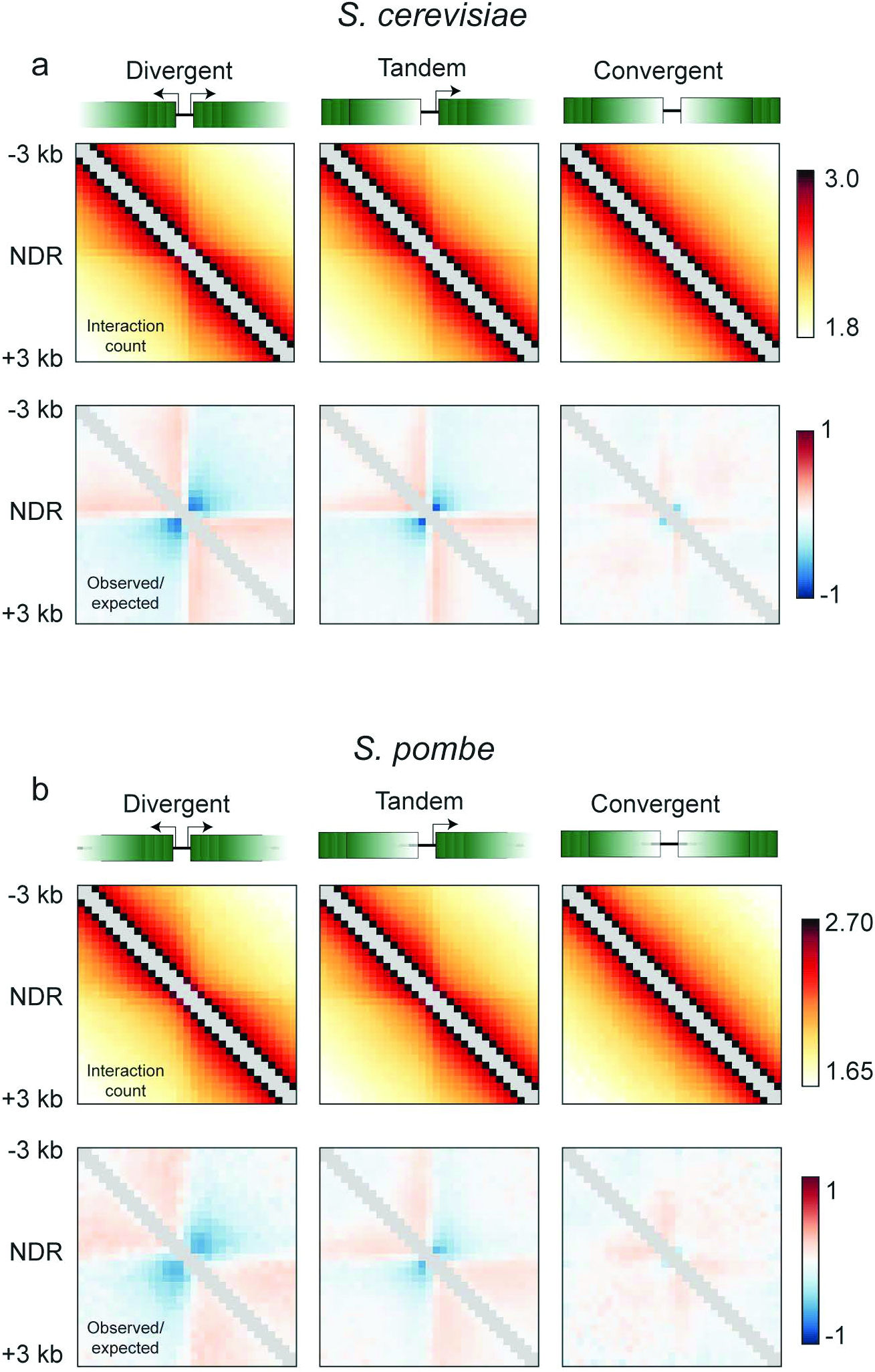
Comparison of chromosome folding in *S. cerevisiae* and *S. pombe*. (**a-b**) For each species, intergenic regions were separated into those falling between pairs of genes oriented divergently, in tandem, or convergently, as indicated, and were aligned according to the midpoint of the nearest respective intergenic region. Data from DSG pellet libraries from *S. cerevisiae* (**a**) or *S. pombe* (**b**) are averaged for all genes in each category. Top panels show interaction counts (normalized to library read depth for coverage-corrected 200 bp-binned maps) for the three classes of intergenic region. Bottom panels show the same data, additionally corrected for the decay in interaction frequency with increasing distance, and expressed as the log2 ratio of observed interactions divided by expected interactions for a given genomic distance.

We conclude that broadly similar principles underlie chromosome folding behavior in these distantly-related fungi, with modest quantitative differences in chromosome structure that could potentially result from the interspecies differences in aspects of genomic structure including chromosome length, gene length, intron abundance, location of rDNA clusters, and nucleosome repeat length.

## DISCUSSION

Here, we present an improved protocol for nucleosome-resolution mapping of chromosome folding, termed Micro-C XL. The primary technical improvements detailed here are 1) the use of additional “long-range” crosslinkers to supplement formaldehyde crosslinking, and 2) fractionation of relatively insoluble chromatin prior to nucleosome ligation and subsequent library construction. Contrary to our initial expectations, the dramatic improvement seen in apparent capture of long-range interactions using these protocols likely results not from the ability of long-range crosslinkers to bridge interacting genomic loci associated with proteins that are more than 3 Å away from one another, but rather from a decrease in the noise caused by soluble nucleosomes encountering one another in solution during the ligation reaction and causing artefactual “interactions” between unlinked nucleosomes (**Fig. S3, Table S1**). This hypothesis is based on the fact that DSG- and EGS-based Micro-C maps are extremely similar despite their substantial difference in crosslinking distance, as well as the finding that isolation of soluble chromatin results in greatly increased noise in Micro-C maps (**Fig. 3**). In addition, we note that chromatin fragments generated by restriction enzymes in typical Hi-C protocols are significantly larger than mononucleosomes, increasing the number of crosslinking opportunities per fragment and thus presumably restricting their diffusion and resultant ability to generate artefactual ligation products. We propose that this difference in fragment size/mobility accounts for the increased noise seen previously in Micro-C relative to standard Hi-C protocols. Further supporting this idea, we find that the improved protocol strongly reduces the incidence of artefactual ligation products between the nuclear genome and the mitochondrial genome, relative to the standard Micro-C protocol (**Fig. S14**). We note this is in general agreement with prior comparisons of in-solution versus both pellet and “in situ” Hi-C protocols^28, 38, 39^. Together, these considerations support the idea that the use of long crosslinkers and isolation of insoluble chromatin may be important to prevent mononucleosomes from freely diffusing prior to ligation and introducing noise into Micro-C measurements. Still, this does not rule out the additional possibility that in some cases our long crosslinkers capture nearby genomic loci for which the closest crosslinkable proteins are not in immediate physical proximity, and indeed both of these features may contribute to the improvement in data quality seen in Micro-C XL.

### Chromosome structure

Validated via comparisons to prior data, our method provides insight into yeast genome folding at all length scales of interest. At larger scales, the Rabl configuration of chromosomes is seen as clustering of centromeres, and interactions between the telomeres of chromosome arms of similar length. Centromeric chromatin also shows a characteristic “X” shape resulting from the two arms of the chromosome both statistically leading away from the centromere together for some ∼20 kb, with centromeres otherwise being relatively isolated from chromosome arms. At higher resolution, genes in both budding and fission yeast are organized into chromosomally-interacting domains (CIDs), typically spanning 1-5 genes, that are in some ways similar to the “topologically-associating domains” described in a multitude of other model organisms. Boundaries between CIDs occur at active promoters, highly-expressed genes, and tRNA genes (**Fig. S15**). In both budding and fission yeast, genomic regions surrounding cohesin-associated loci are relatively insulated from physically interacting with one another (**Fig. S16**). However, this insulation is stronger and persists over greater genomic distances in *S. pombe* relative to *S. cerevisiae*, pointing towards important differences in the role of cohesin, potentially in a cell-cycle dependent fashion, between the species. Taken together, these analyses highlight the ability of Micro-C XL to assay chromosome folding across all scales, as well as its broad applicability and future utility in comparative genomics.

### Applications to other biological systems

We finally turn to considerations of the sequencing depth required for applications of Micro-C XL in organisms with larger genomes. As reviewed in Lajoie *et al*^29^, the fundamental genomic resolution of a chromosome capture dataset is set by the frequency at which the genome is fragmented prior to capture of physical interactions by ligation; beyond this lower bound to resolution, the effective genomic resolution is further influenced by sequencing depth and library complexity. The proportion of molecular byproducts in a library additionally influences the amount of sequencing required to achieve a given coverage per fragment. Given that Micro-C XL does not display a preponderance of molecular byproducts (**Fig. S14**), the sequencing depth required to achieve a given genomic resolution should be similar to a Hi-C protocol. Nevertheless, Micro-C XL has the capacity to analyze chromatin interactions at genomic distances smaller than currently available Hi-C protocols. Indeed, the highest-resolution studies performed to date in mammals^28^ utilize restriction enzymes with 4 bp target sequences to yield average fragment lengths of ∼256 bp, although due to the heterogeneous distribution of restriction sites across the genome lower-resolution (∼1 or 5 kb) binning approaches must be used in analysis of such datasets. In comparison to any individual 4-cutter, MNase digestion of chromatin to mononucleosomes results in at most ∼75% more genomic fragments (depending on the nucleosome repeat length in the tissue of interest), which in turn increases the fundamental genomic *resolution* of Micro-C by substantially more than ∼1.75-fold, thanks to the more even spacing of the resulting fragments.

In addition, beyond binning Micro-C data to mimic lower-resolution Hi-C, it is important to note that a wide variety of biological questions can be addressed – at high resolution – by Micro-C at much lower sequencing depth. First of all, the strength of the Micro-C protocol is its ability to interrogate chromatin fiber structure at ∼150-1000 bp resolution – there is little reason to carry out Micro-C to investigate >1 MB chromatin domains^19^. A key measure in this regard 40 is the decay of interaction frequency with increasing distances (see, e.g., **Fig. 1d** or **Fig. 3a**), which is an averaged measure across the entire genome and is thus extremely robust to undersequencing (**Fig. S17**). We anticipate that very low coverage (below 1-2 million reads) Micro-C XL maps in mammals will thus allow robust comparison of average chromatin fiber folding for, say, Polycomb-repressed genes, or for exons vs. introns, etc. In addition to using such computational averaging methods to make use of multiple instances of any given annotation, the molecular complexity of the sequencing library can also be experimentally reduced. This is commonly done in sequence-capture RNA-Seq protocols in cancer exome studies, and more recently such methods have been applied to 3C methods based either on protein capture (eg, Pol2 IP) or on capture of specific “bait” sequences^41^. The reduction in read depth required for such methods naturally depends on the distribution and abundance of the feature to be captured, but many proteins of interest – CTCF, cohesin, TFIIB, and others – are sparsely distributed enough to enable >100-fold reductions in the sequencing depth required for highresolution Hi-C studies. These and other considerations^29, 41^ must be a part of any experimental design for a 3C-based study.

### Conclusion

Here, we describe a modified protocol for genome-wide analysis of 3D chromatin structure that captures aspects of chromosome folding at all scales from mononucleosome resolution up to interactions between different chromosomes. This protocol, Micro-C XL, should find broad utility in a multitude of biological systems.

## METHODS

### Yeast strains and culture conditions

All experiments reported here were carried out with either *S. cerevisiae* strain BY4741 or *S. pombe* strain 972 h-. BY4741 cultures were grown in YPD media at 30°C, while *S. pombe* cells were grown at 30°C in “Compromise Media”^42^, consisting of Yeast extract (1.5%), Peptone (1%), Dextrose (2%), SC Amino Acid mix (Sunrise Science) 2 grams per liter, Adenine 100 mg/L, Tryptophan 100 mg/L, and Uracil 100 mg/L.

### Fixation conditions

Midlog yeast cultures were crosslinked with either 1% or 3% final concentration of formaldehyde (Sigma) for 15 minutes at 30°C, then quenched with 125 mM glycine for 5 min at room temperature. Yeast were then spheroplasted as previously described^42, 43^. For DSG and EGS (Thermo Fisher Scientific) crosslinking studies, spheroplasts were resuspended in a 3 mM final concentration of the crosslinker of interest in PBS, and crosslinked for 40 min at 30°C, then quenched with 125 mM glycine for 5 min at room temperature. Note that we use the protocol “Micro-C XL” for DSG and EGS-based protocols interchangeably, as the data from these protocols are nearly indistinguishable.

### Separation of soluble and insoluble chromatin

Crosslinked chromatin was digested with micrococcal nuclease (MNase, Worthington) to yield > 95% mononucleosomes. After inhibition of MNase with 2 mM EGTA at 65°C, fragmented lysate was in some cases used directly for the standard Micro-C protocol, or in some experiments (**Figs. 3-5**) was separated into supernatant and pellet portions. Here, MNase-digested lysate was spun at 10,000 g for 5 minutes, and Micro-C was separately performed on supernatant or on the pellet fraction.

### Micro-C protocol

The termini of nucleosomal DNA were dephosphorylated by Shrimp Alkaline Phosphatase and then subjected to T4 DNA polymerase for end repairing and biotin labeling by supplementing with biotin-dCTP, biotin-dATP, dTTP, and dGTP. Crosslinked chromatin was diluted to 10 ml and treated with T4 DNA ligase. After heat inactivation, chromatin was concentrated to 250 µl in an Amicon 30k spin column and treated with 100 U exonuclease III for 5 min to eliminate biotinylated ends of unligated DNA. Proteinase K was then added and incubated for 65°C overnight. DNA was purified by PCI extraction and ethanol precipitation, treated with RNase A, and ∼250–350 bp DNA was gel-purified. Purified DNA was treated with End-it, subject to A-tailing with Exo-Klenow, and ligated to Illumina adapters. Adapter-ligated DNA was purified with streptavidin beads to isolate ligated Micro-C products away from undigested dinucleosomal DNA. Streptavidin beads were then subject to ∼10–12 cycles of PCR using Illumina paired-end primers. Amplified library was purified and subject to Illumina NextSeq paired end sequencing. Detailed methods are described in **Supplemental Text**.

### Computational analysis of Micro-C interactions

MicroC data was mapped to the sacCer3 genome using Bowtie 2.1.0 as described in^44^ using the hiclib library for python, publicly available at **https://bitbucket.org/mirnylab/hiclib**, with virtual 100bp fragments. To obtain corrected contact maps, genomic coverage was calculated by summing the total number of interactions per bin. Low coverage bins were then excluded from further analysis using a MAD-max (maximum allowed median absolute deviation) filter on genomic coverage, set to 9 median absolute deviations. Following this filtering, standalone bins were removed (ie. regions where both neighboring bins did not pass filters), and the resulting maps were then iteratively corrected to equalize genomic coverage^44^. Observed/expected contact maps were obtained by additionally dividing out the dependence on genomic distance, calculated empirically as the mean number of contacts at each genomic separation, using a sliding window with linearly increasing size, as previously described^45^. Log-log plots of contact probability *P(s)* versus distance were calculated using log-spaced bins with a constant step size. For average plots around genomic features, gene positions and orientations, centromere positions, and tRNA positions were obtained from the SGD (http://www.yeastgenome.org/).

## ACKNOWLEDGEMENTS

We thank L. Mirny and members of the Rando lab for insightful discussions. Work was supported in part by NIH grant GM079205 to OJR. GF and AG were supported by R01 GM114190 and U54 DK107980 of the 4D Nucleome Program. T-HSH is an HHMI international student research fellow.

## SUPPLEMENTARY FIGURES

**Fig. S1.**
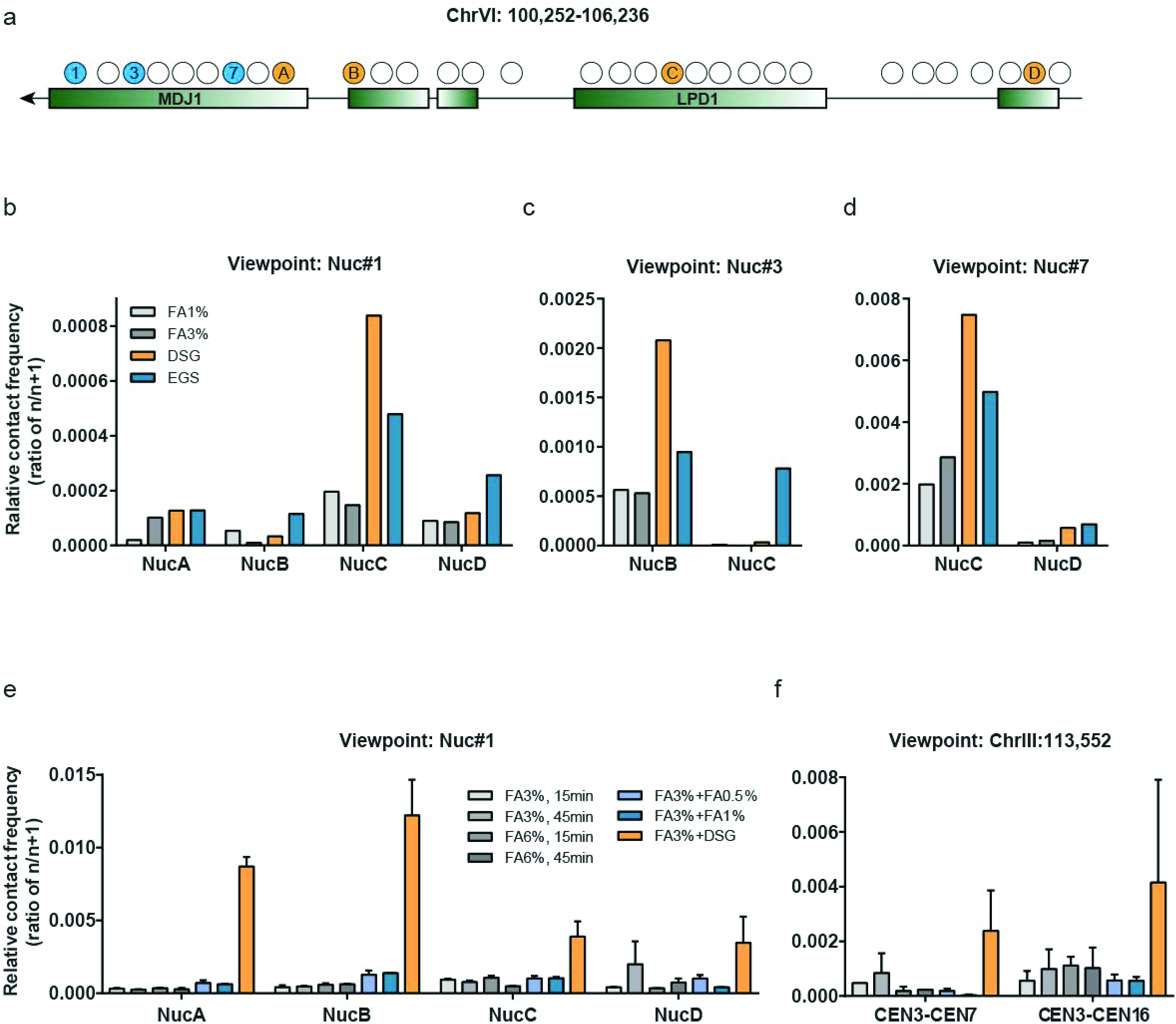
Q-PCR analysis of effects of long crosslinkers on Micro-C protocol. Budding yeast were crosslinked with formaldehyde, permeabilized, and then treated with one of several alternative crosslinkers. Mononucleosomal DNA was then processed using the Micro-C protocol, and ligated DNA was subject to q-PCR using primer pairs designed against a variety of nucleosomes surrounding the *MDJ1* contact domain. (**a**) Schematic of primer locations for q-PCR analyses of the ∼6 kb region surrounding *MDJ1*. Forward primers located at nucleosomes 1, 3, and 7 are indicated in blue, while locations of reverse primers are shown in orange. (**b-d**) q-PCR data for the indicated forward and reverse primer pairs, normalized to the q-PCR signal obtained to the abundance of ligation products between the upstream nucleosome in question and its immediate downstream neighbor (e.g., for panel (**c**) data are normalized to the pairwise interaction between *MDJ1* nucleosomes number 3 and 4). Data are shown for 1% FA, 3% FA, 3% FA + DSG, and 3%FA + EGS, as indicated. (**e**) As in (**b**), but showing data for additional crosslinking conditions including higher FA concentrations, longer FA crosslinking, and two-step FA crosslinking in which a second FA incubation is carried out after spheroplasting. This last protocol mimics the use of DSG or EGS after spheroplasting in the Micro-C XL protocol. (**f**) q-PCR data for interactions between CEN3 and two other centromeres, showing data for the same protocols detailed in (**e**).

**Fig. S2.**
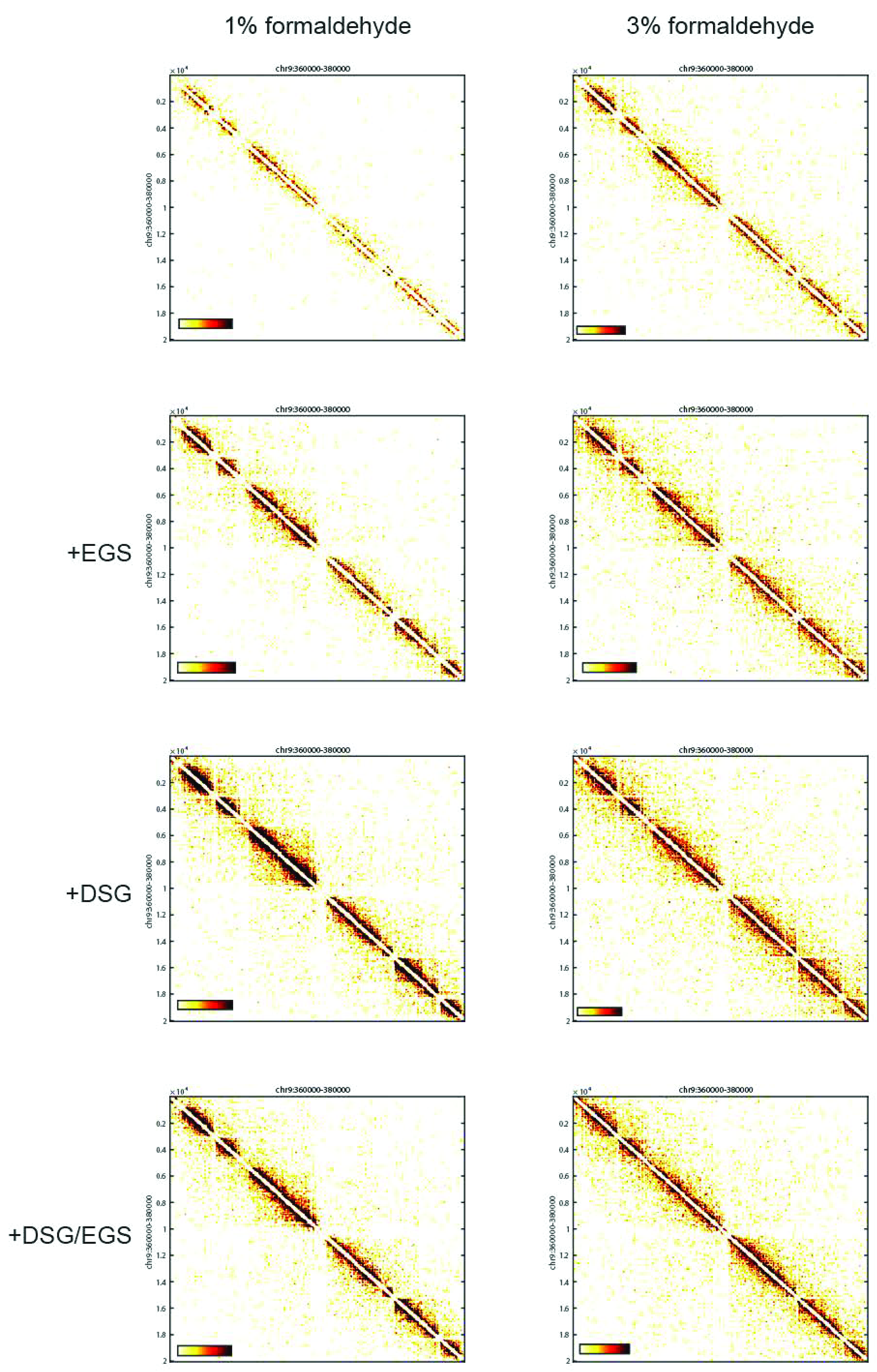
Comparison of crosslinking protocols for a typical 20 kb region. Micro-C data are shown for chrIX: 360,000-380,000 for eight different crosslinking conditions, as indicated. The raw matrix was only normalized to sequencing depth as described in **Fig. 1c** and Hsieh *et al*^17^, and interactions were counted in single bp resolution without binning. Improved capture of contact domains associated with individual genes is readily apparent here for protocols incorporating DSG or EGS.

**Fig. S3.**
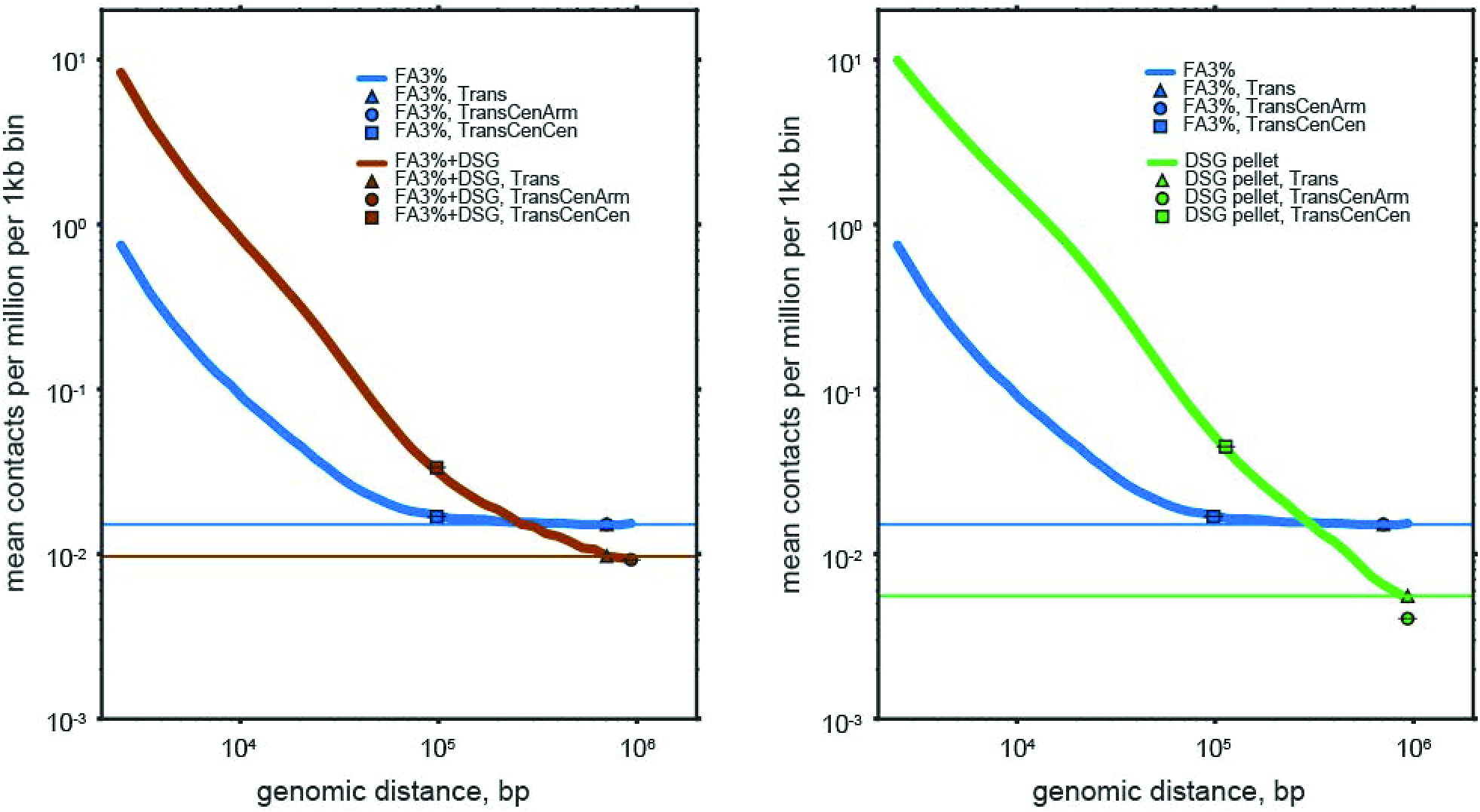
Addition of long crosslinkers reduces the “noise floor” relative to FA-only Micro-C maps. Intra-arm contact probability, *P(s)*, as a function of genomic distance, *s*, was calculated from 1kb corrected contact maps as in^44^, using 50 logarithmically spaced bins from 1kb to 1Mb. Horizontal line marks average trans (between-chromosome) contact frequency. Markers respectively indicate average trans Cen-vs-Cen and trans Cen-vs-Arm contact frequencies, defining each centromere with a +/-20kb genomic window. Note that *P(s)* flattens out at the average trans contact frequency in the FA3-only dataset, as would be result from an adding a constant frequency of interaction between any two loci. Additionally, while trans-cen-arm and trans-average are similarly strong in the FA3-only and FA3-DSG datasets, in the DSG-pellet dataset, the avoidance of the centromere from arm regions is clearly seen.

**Fig. S4.**
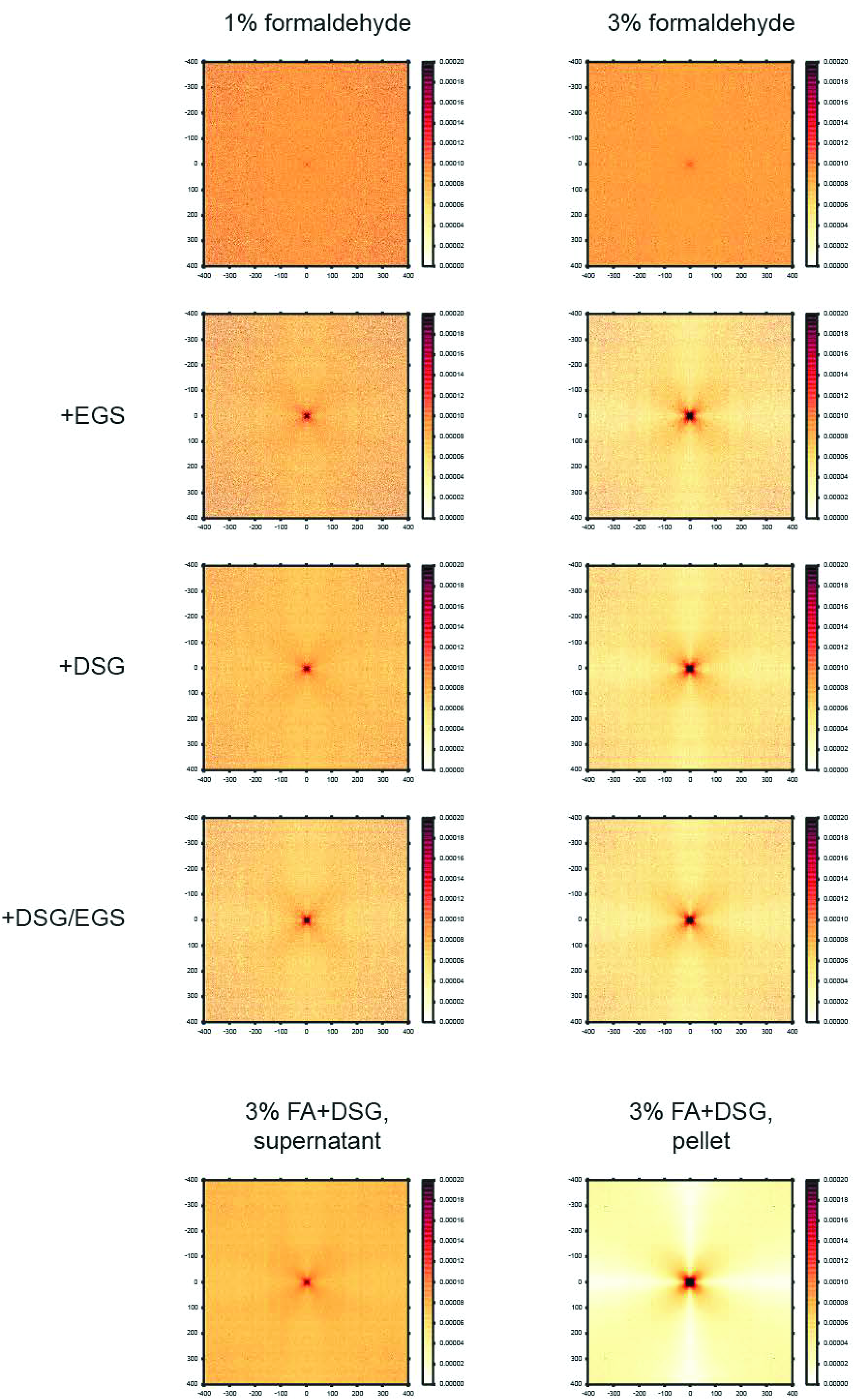
Centromere clustering revealed by alternative crosslinkers. Average interaction map for all possible pairs of CEN-CEN interactions for the indicated Micro-C protocols. Top eight panels show standard Micro-C protocol performed using the indicated crosslinking conditions, while bottom two panels show data for Micro-C performed following separation of relatively soluble and insoluble MNase-digested chromatin by centrifugation prior to ligation. 1kb binned contact maps were corrected for genomic coverage and normalized such that the total coverage of each 1kb region summed to 1.

**Fig. S5.**
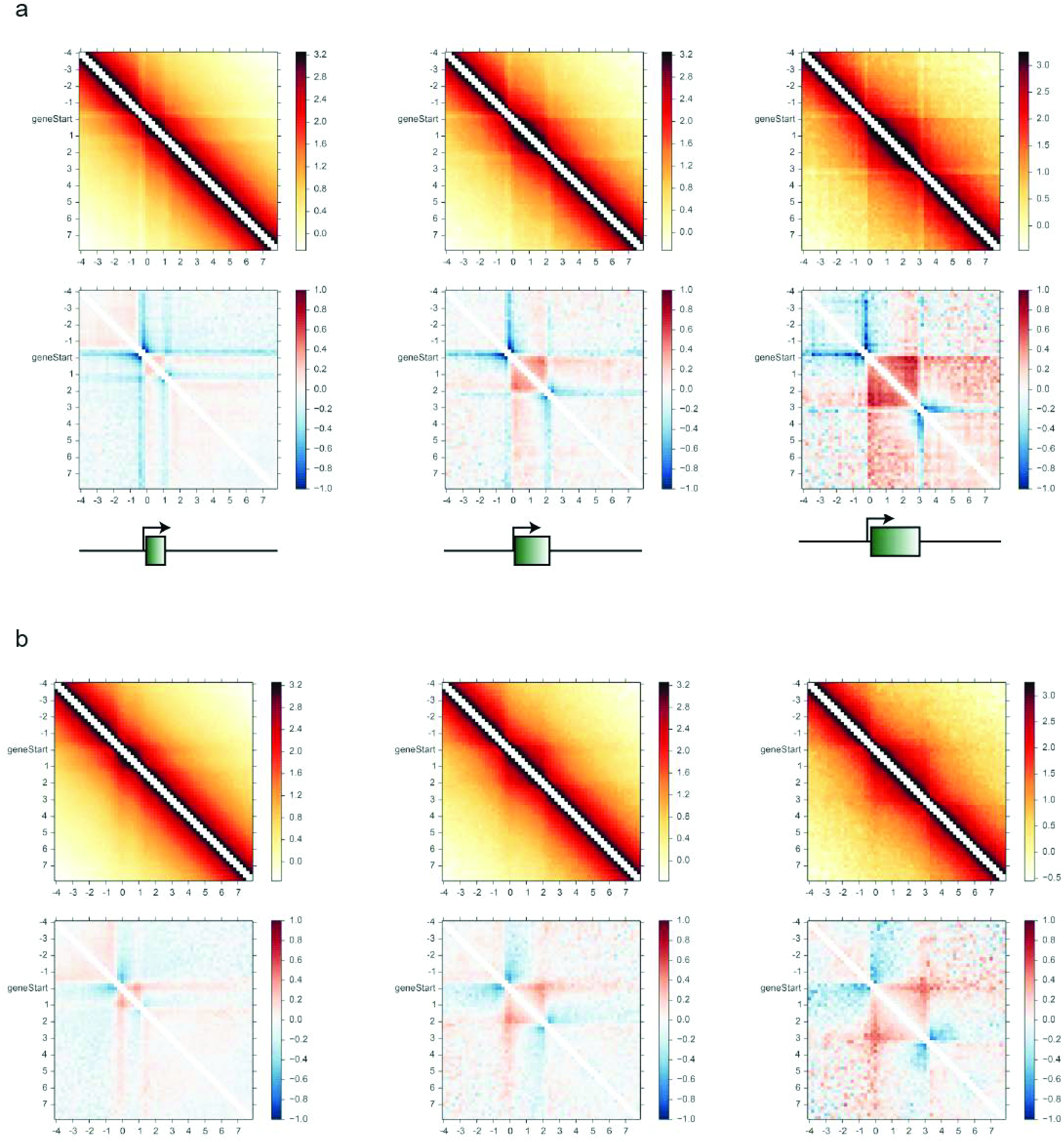
Effects of normalization on Micro-C metagenes. (**a**) Metagene maps for *S. cerevisiae* DSG pellet dataset, binned to 200bp resolution and normalized only for sequencing depth. All genes of length 1-1.2 kb, 2-2.2 kb, and 3-3.2 kb, as indicated, were identified and aligned by their 5’ ends. The narrow range of gene lengths was chosen to assist in visualization of a discrete 3’ gene end in these plots. Top panels show log10 averaged interaction counts, normalized only for library read depth. CID structure is evident in these panels as a region of increased contacts bounded at both the 5’ and the 3’ ends of genes. Note that interactions within each box decay smoothly with increasing distance from the diagonal, indicating that interactions between gene ends are at most a minority subpopulation of gene folding conformations. Bottom panels show the same data, after additionally controlling for the global decay in interaction frequency with increasing genomic distance. Data are shown as log2 of the observed interactions divided by the interaction count expected based on genomic distance. This correction reveals a far clearer view of CID structure, with clear blue boundaries delimiting the red contact domain associated with the gene. (**b**) As in (**a**), but data are additionally normalized by matrix balancing, which corrects for nucleosome occupancy (observed as variation in the total coverage per bin in Micro-C maps). Visually this removes the faint “stripes” in the raw data (top panels) associated with nucleosome-depleted promoter regions. Following this normalization a subtle enrichment of interactions can be observed for the +1/+N nucleosome interaction in the observed/expected visualization (bottom row). Nevertheless, we note that even uniform squares on an observed map can have apparent corner peaks after dividing by an expected map where contact frequency decreases with genomic distance. Moreover, regardless of normalization scheme, we do not observe an uptick in contact probability between the +1/+N nucleosomes in observed maps for any of the assayed gene lengths, the signature of a gene loop. For these reasons, micro-C data argues for the prevalence of gene- and multi-gene-wide crumpling, rather than specific +1/+N gene loops.

**Fig. S6.**
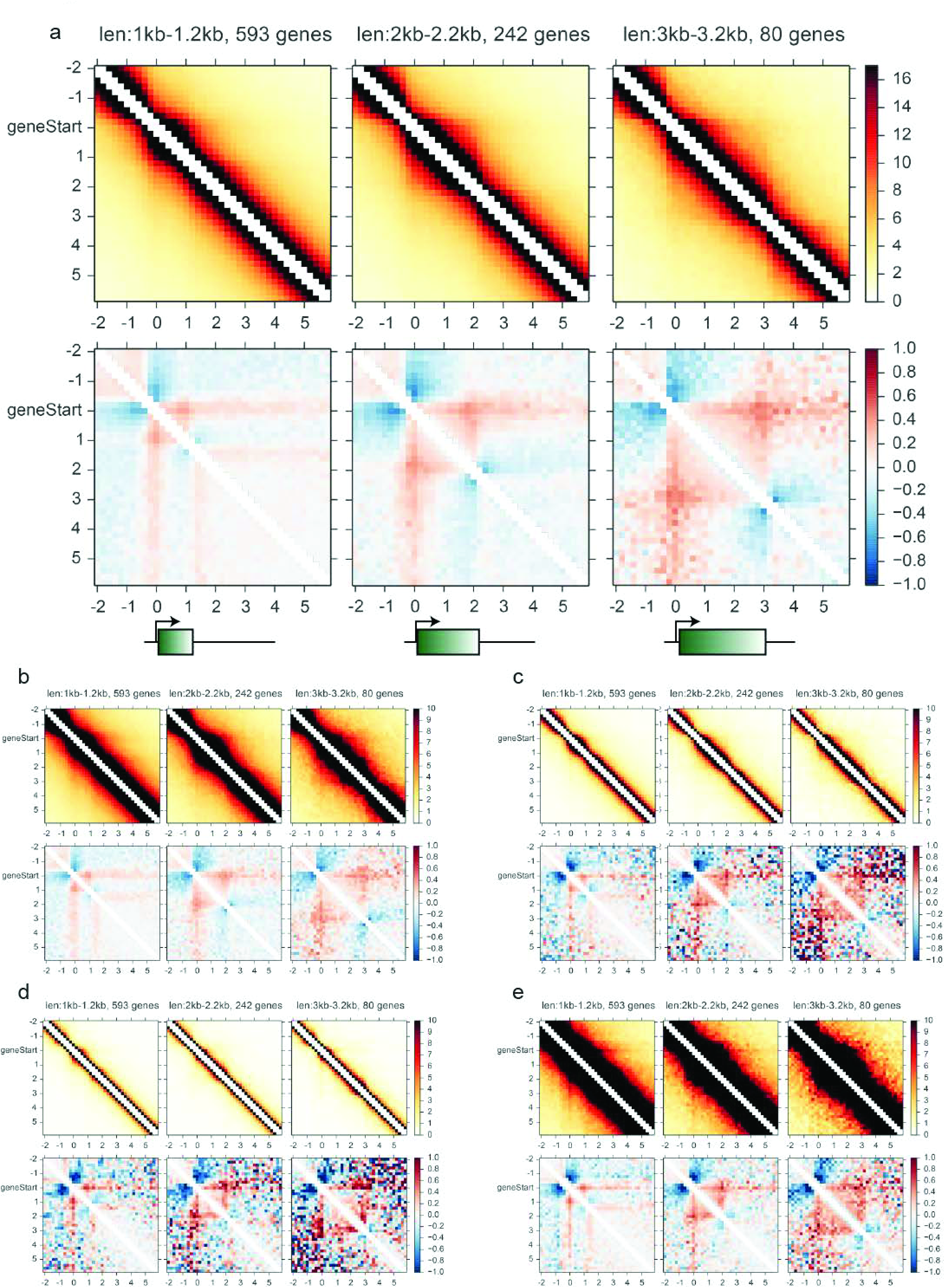
Metagene analysis of genes grouped by length is robust to protocol variation. Metagene visualization as in **S5** for (**a**) EGS pellet, (**b**) EGS supernatant, (**c**) 3% FA, and (**d**) 3% FA + DSG, Micro-C libraries.

**Fig. S7.**
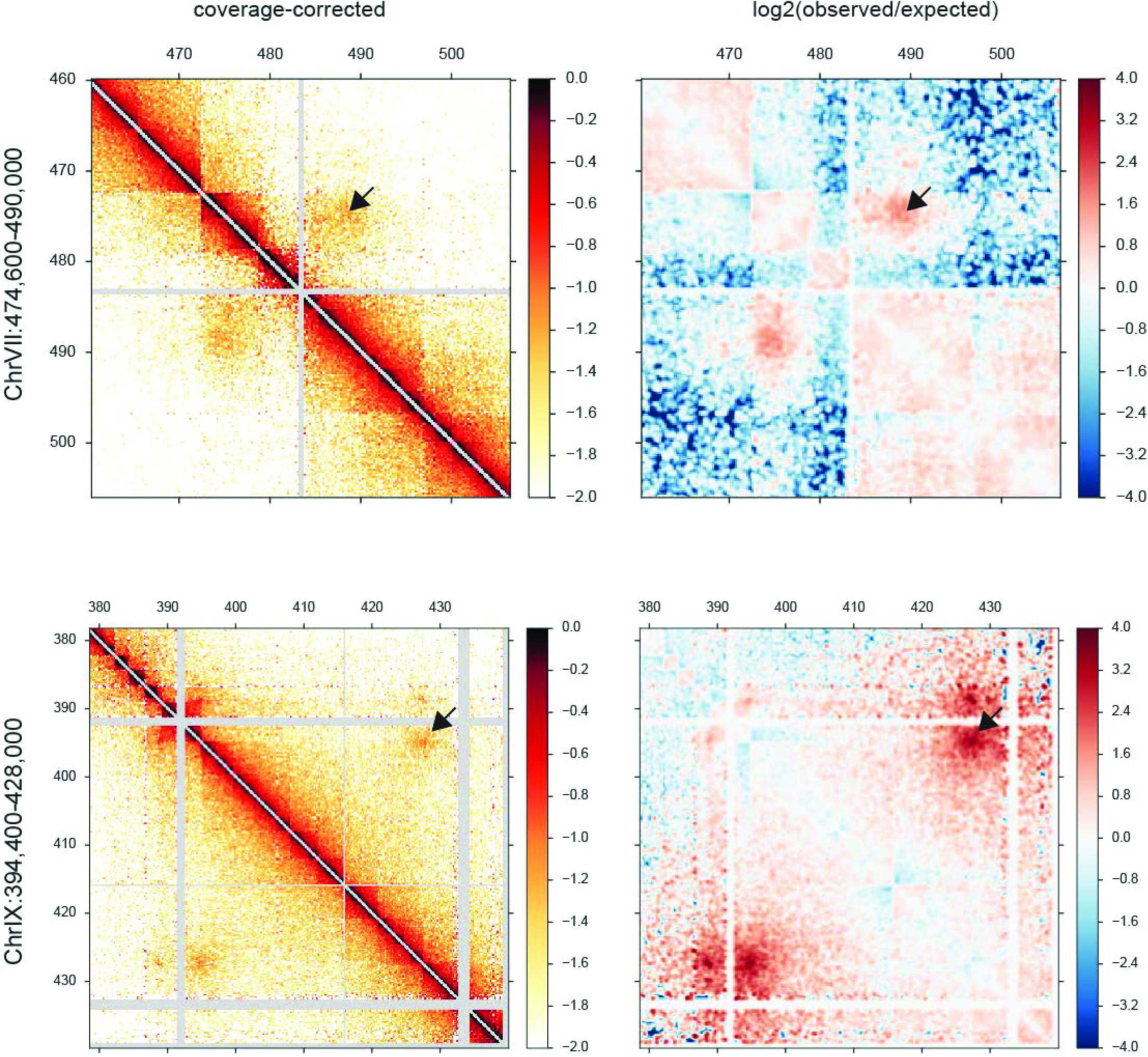
Example of two potential looping interactions. Data for the indicated loci are shown using 200bp binned maps, with left panel showing interaction counts for DSG pellet libraries normalized only for sequencing depth, and right panel showing smoothed observed/expected ratio. Clear in both cases is a possible chromosomal looping interaction. Both loops are observed much more strongly in DSG pellet libraries than in supernatant libraries (not shown).

**Fig. S8.**
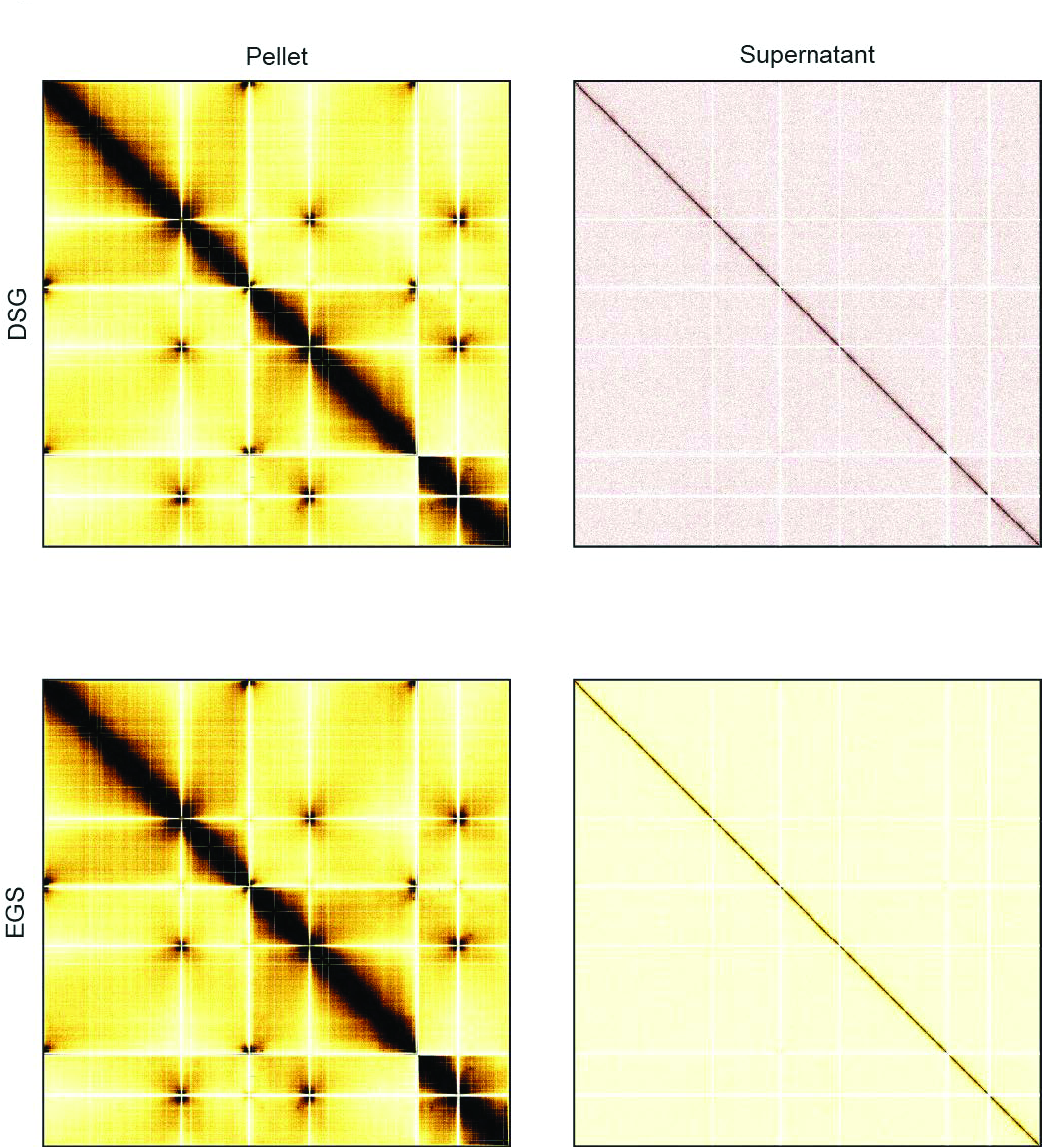
Isolation of insoluble chromatin improves signal to noise for *S. pombe* Micro-C maps. Whole genome contact Micro-C XL maps were generated using FA+DSG or FA+EGS as crosslinkers, and were generated from soluble or insoluble material, as indicated, for the fission yeast *S. pombe*. Maps are normalized for sequencing depth only.

**Fig. S9.**
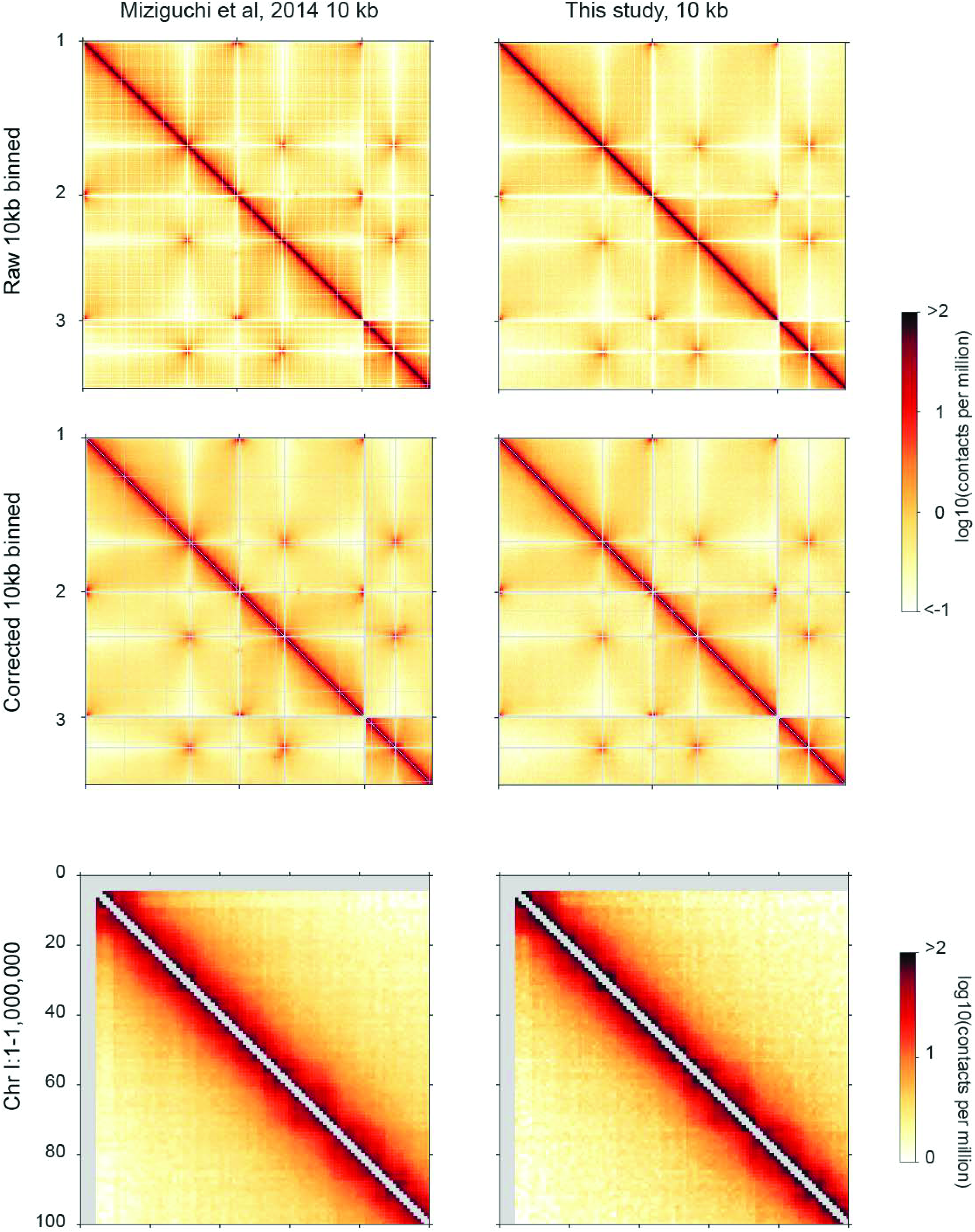
Comparison of Micro-C XL and published Hi-C maps for *S. pombe* at 10 kb resolution. In each row, left panels show data from Mizuguchi *et al*^24^, while right panels are from this study, binned at 10 kb resolution. Top two rows show data for the entire genome, while bottom row shows a 1 MB zoom-in. Overall Micro-C XL maps are highly-correlated with published results for this species (Spearman’s r = 0.77 for corrected 10 kb resolution maps, comparable with the r = 0.77 correlation between DSG and EGS pellet maps for *S. cerevisiae*), with both maps showing ∼100 kb chromatin contact domains previously referred to as chromatin “globules”.

**Fig. S10.**
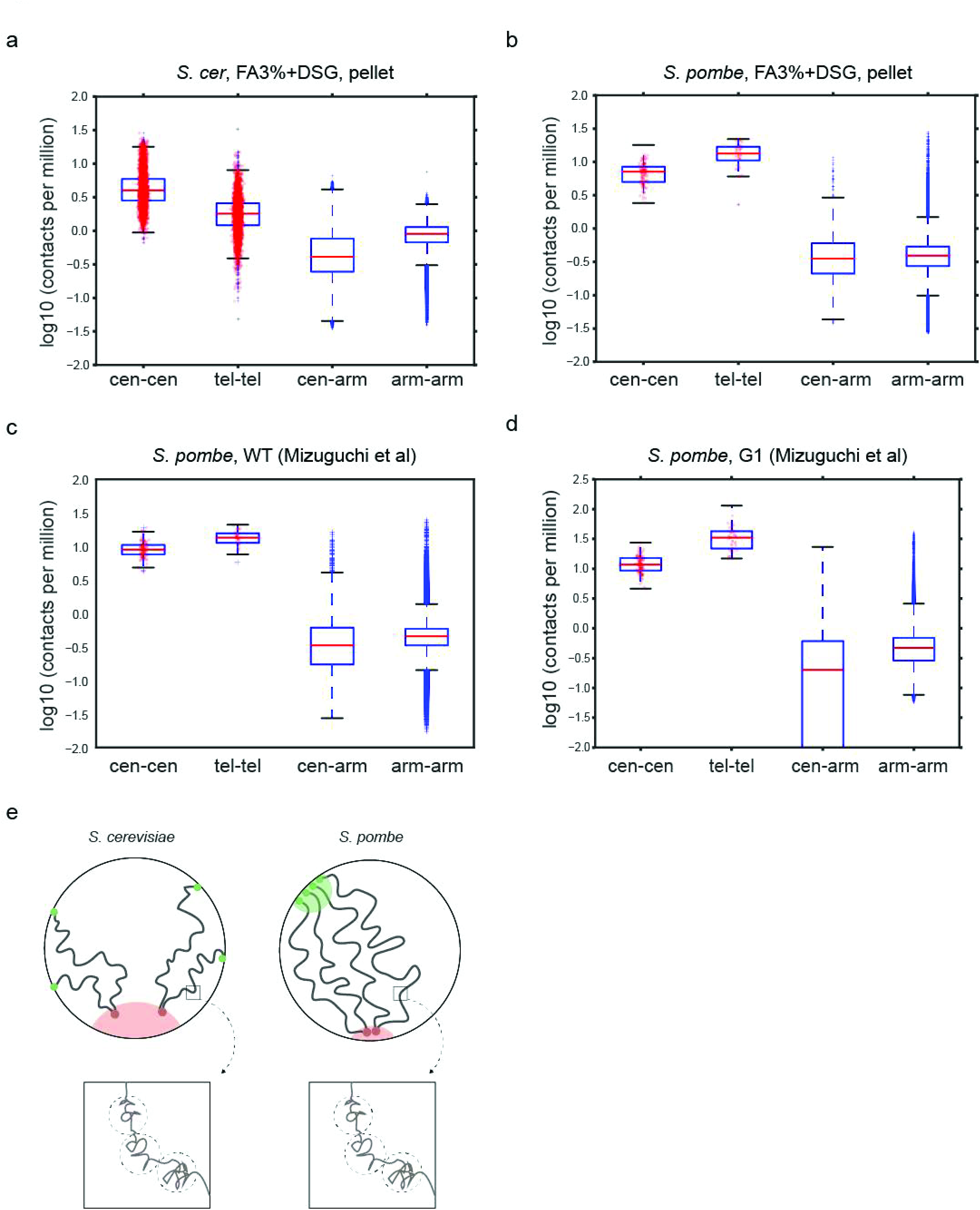
Comparison of global folding behavior between two yeast species. (**a-d**) Boxplots showing fraction of contacts (normalized as parts per million read pairs) for interactions between centromeres (cen-cen), between telomeres (tel-tel), between centromeres and chromosome arms (cen-arm), and between distal chromosome arms (arm-arm), calculated from coverage-corrected 10kb binned contact maps. Chromosome arms here are defined as sequences more than 20 kb away from either a centromere or telomere. Boxplots here show the median (red line), the first and third quartiles (box), and 1.5* the inner quartile range (whiskers). Red points overlay values for bin-pairs for (cen-cen) and (tel-tel) regions. As in^24^, the 10 most telomere-proximal bin-pairs for non-filtered regions of the heatmap are chosen, or 40 most centromere-proximal bin pairs (as there are 4 arm pairs at each centromere). Data for (**a**) and (**b**) are taken from this study, while (**c**) and (**d**) show data from Mizuguchi *et al* for unsynchronized or G1-arrested *S. pombe* (**c** and **d**, respectively). Note that the enhanced telomere clustering seen in *S. pombe* relative to *S. cerevisiae* is observed both using Micro-C XL (**b**) and Hi-C (**c**) in unsychronized *S. pombe*, and in G1-arrested (**d**) *S. pombe*, indicating that this difference between budding and fission yeast does not result from the substantial difference in their cell cycle dynamics. (**e**) Cartoon models of budding and fission yeast chromosome folding. Fission yeast exhibit a subtle enhancement in centromere clustering, as well as much more substantially-enhanced telomere clustering (left panels), but at the level of individual genes (right panels) fission yeast chromosomes exhibit similar folding properties to budding yeast chromosomes.

**Fig. S11.**
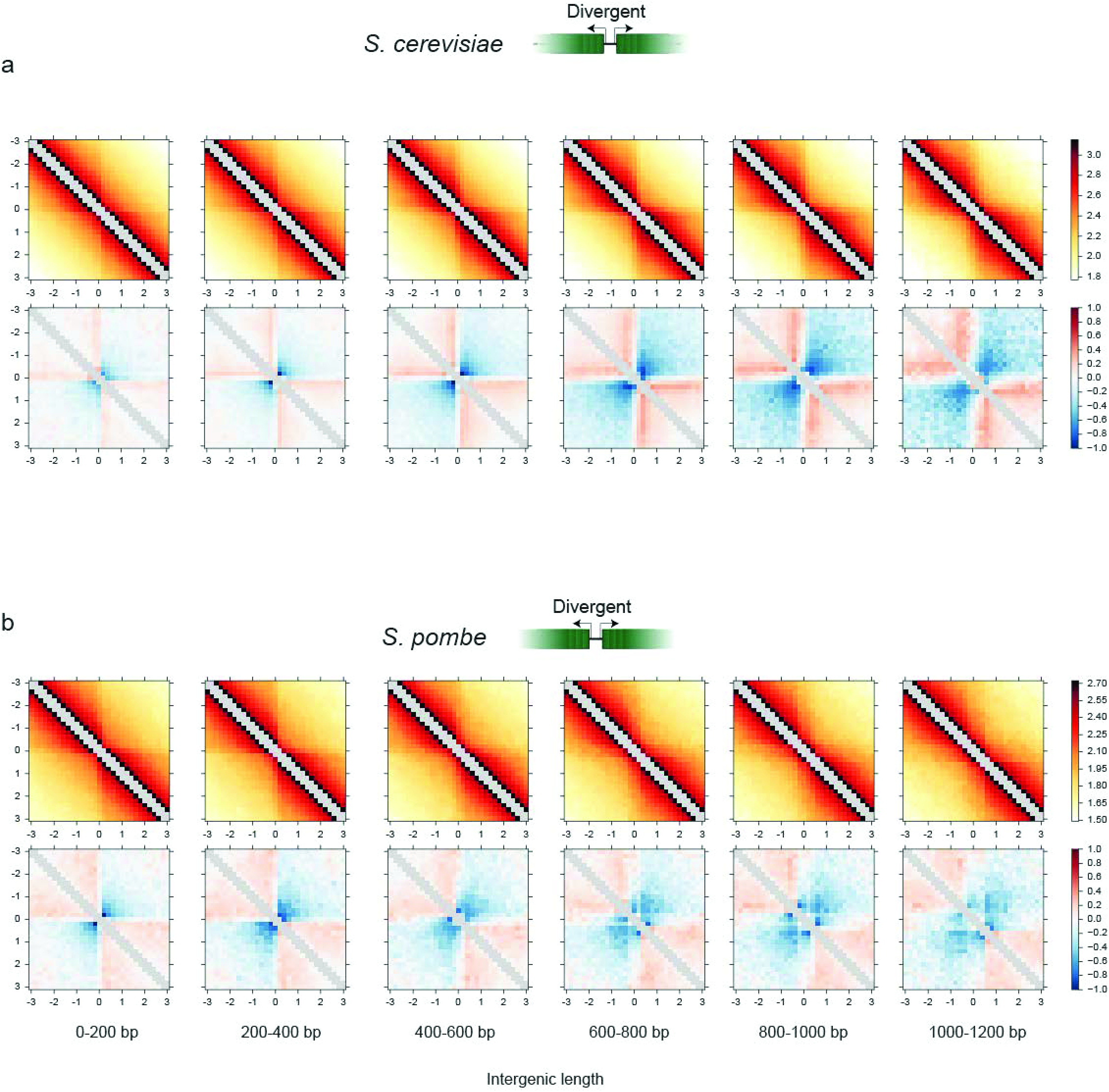
Comparison of boundary activity of promoters in budding and fission yeast. For both species, two rows of panels are shown as in **Figs. S5-6** and **Fig. 5**, with interaction counts in top panels and distance-corrected interaction levels in bottom panels. In both cases data is from DSG pellet maps. Data here are shown for divergently-oriented genes, separated into groups based on the intergenic distance, as shown. In both species divergent promoters act as boundaries, with longer intergenic regions more effectively separating chromatin domains from one another. Budding and fission yeast also exhibit similar behavior at intergenic regions separating tandemly-oriented genes, and separating convergently-transcribed genes (not shown).

**Fig. S12.**
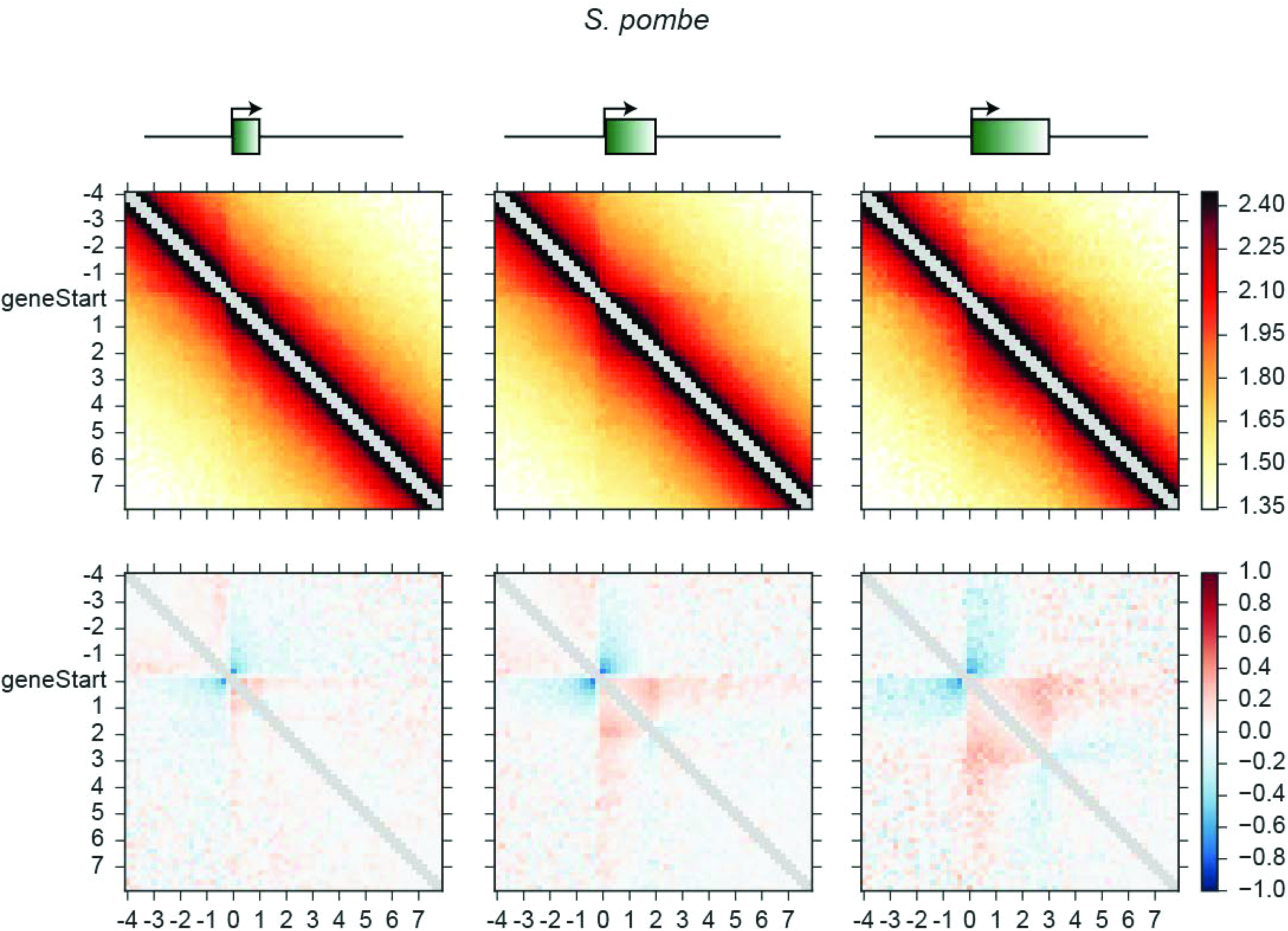
Metagene analysis of *S. pombe* genes. As in **Fig. S6**, for *S. pombe* DSG pellet data. As observed for *S. cerevisiae*, gene loops are not observed in coverage-corrected interaction data, but distance-corrected interactions reveal compacted domains at the gene level.

**Fig. S13.**
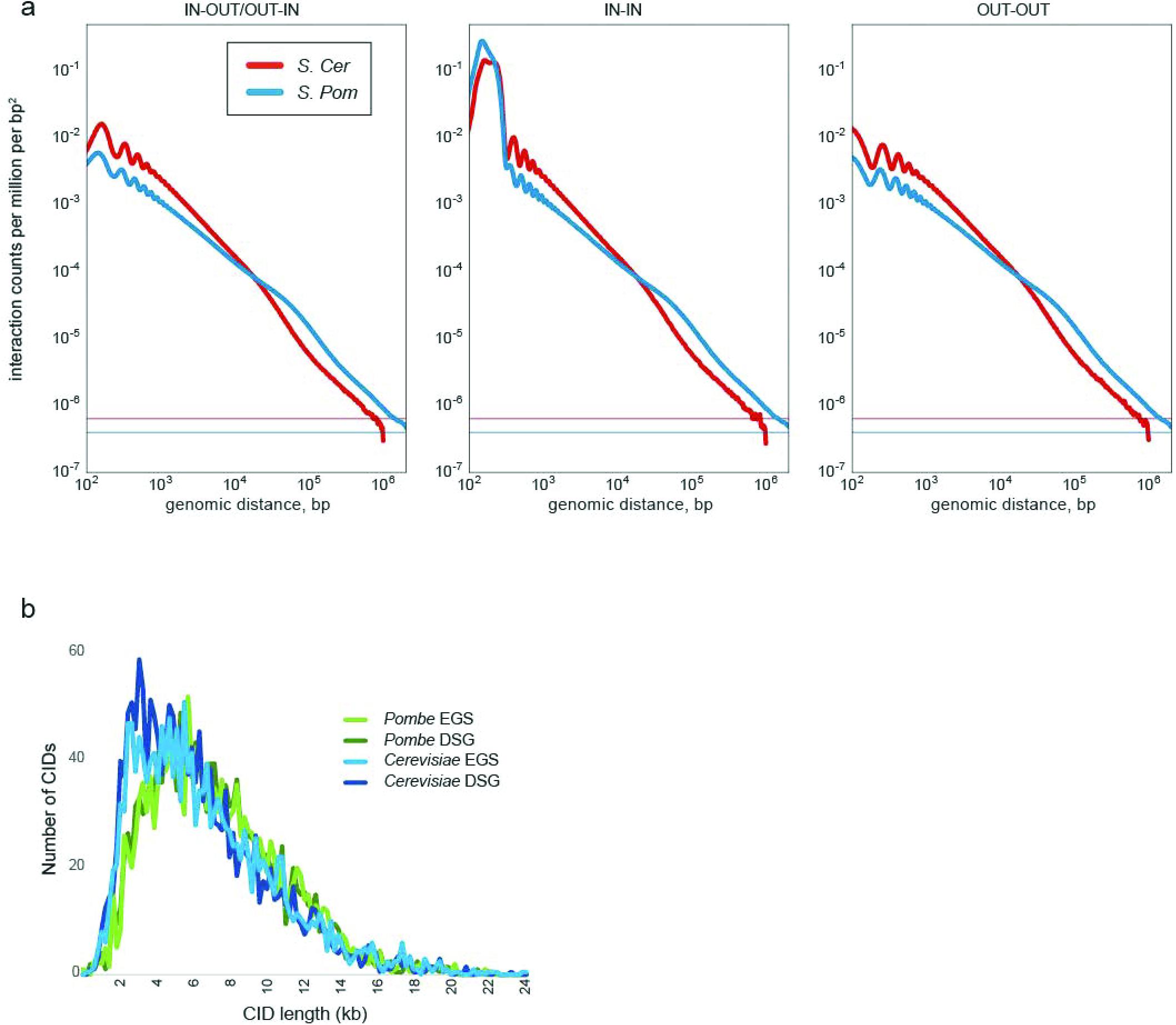
Comparison of *S. pombe* and *S. cerevisiae* genome folding. (**a**) Decay of Micro-C XL interactions with increasing genomic distance. Interactions vs. distance are shown for the indicated read pair orientations for the two species. Subtle differences at short distances are primarily attributable to different nucleosome repeat lengths in these species, while at longer distances we find a subtle signal for *S. pombe* interactions decaying more slowly with increasing distance. Interestingly, we also note an inflection point at ∼70 kb at which interactions in *S. pombe* decay more rapidly – this may reflect the more robust organization of the fission yeast genome into cohesin-delimited “globules”. (**b**) Distribution of contact domain lengths. Boundaries between contact domains were called as described in Hsieh *et al* for *S. cerevisiae* and *S. pombe* Micro-C XL datasets. Plots show the distribution of lengths for boundary-delimited contact domains, which are extremely similar for these two species.

**Fig. S14.**
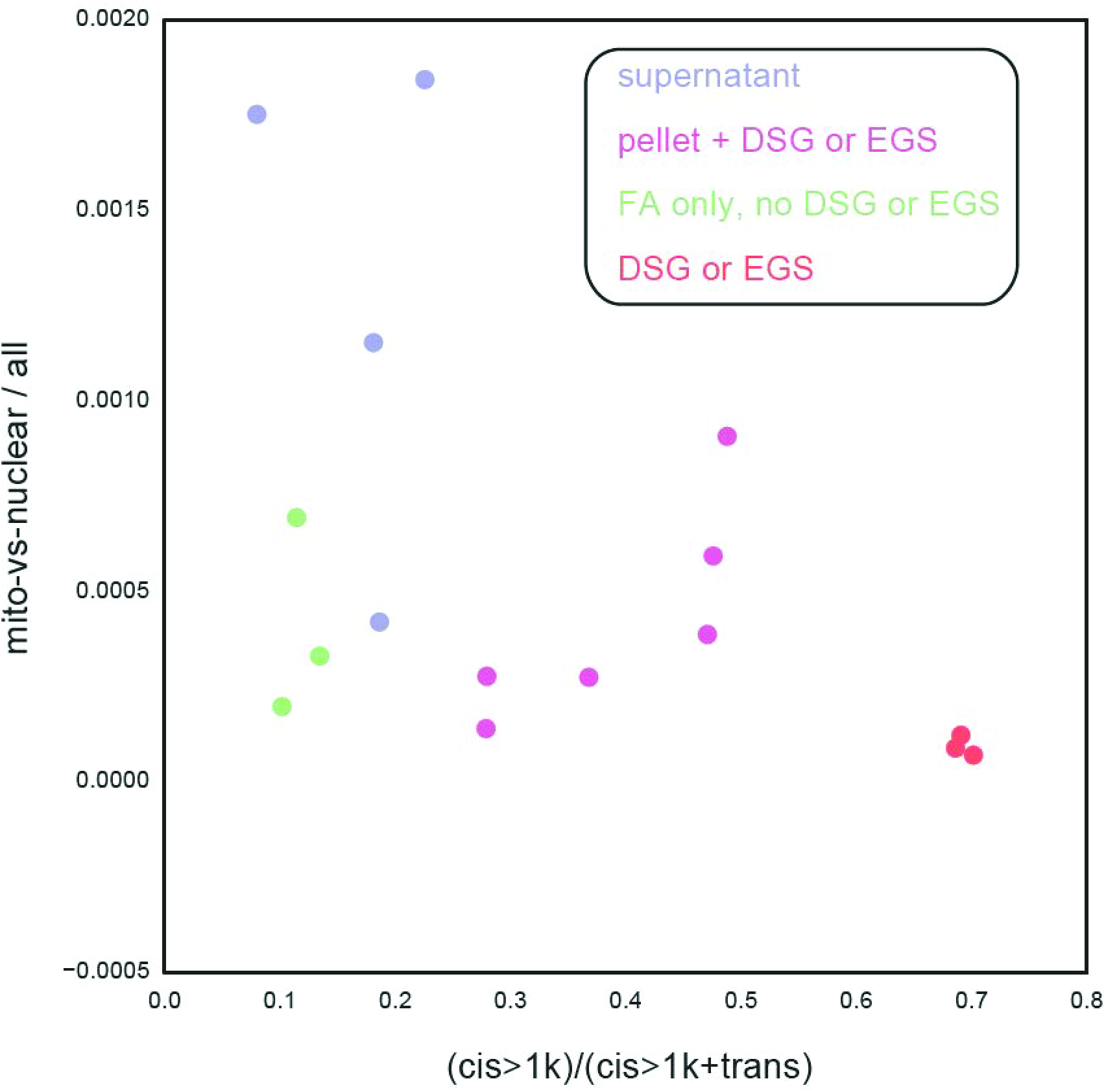
Effects of Micro-C protocols on artefactual interactions. For each Micro-C dataset generated for *S. cerevisiae* in this study, we calculated 1) the fraction of sequencing reads mapping to the mitochondrial genome, and 2) the ratio between those reads reporting on an interaction between two loci on the same chromosome (in cis, >1 kb), and reads reporting on an interaction between chromosomes (in trans). Here, these two values are scatterplotted against one another for all Micro-C datasets. Note that supernatant libraries exhibit a greater frequency of mitochondrial reads relative to other Micro-C libraries, and that pellet libraries exhibit a strong depletion of trans interactions.

**Fig. S15.**
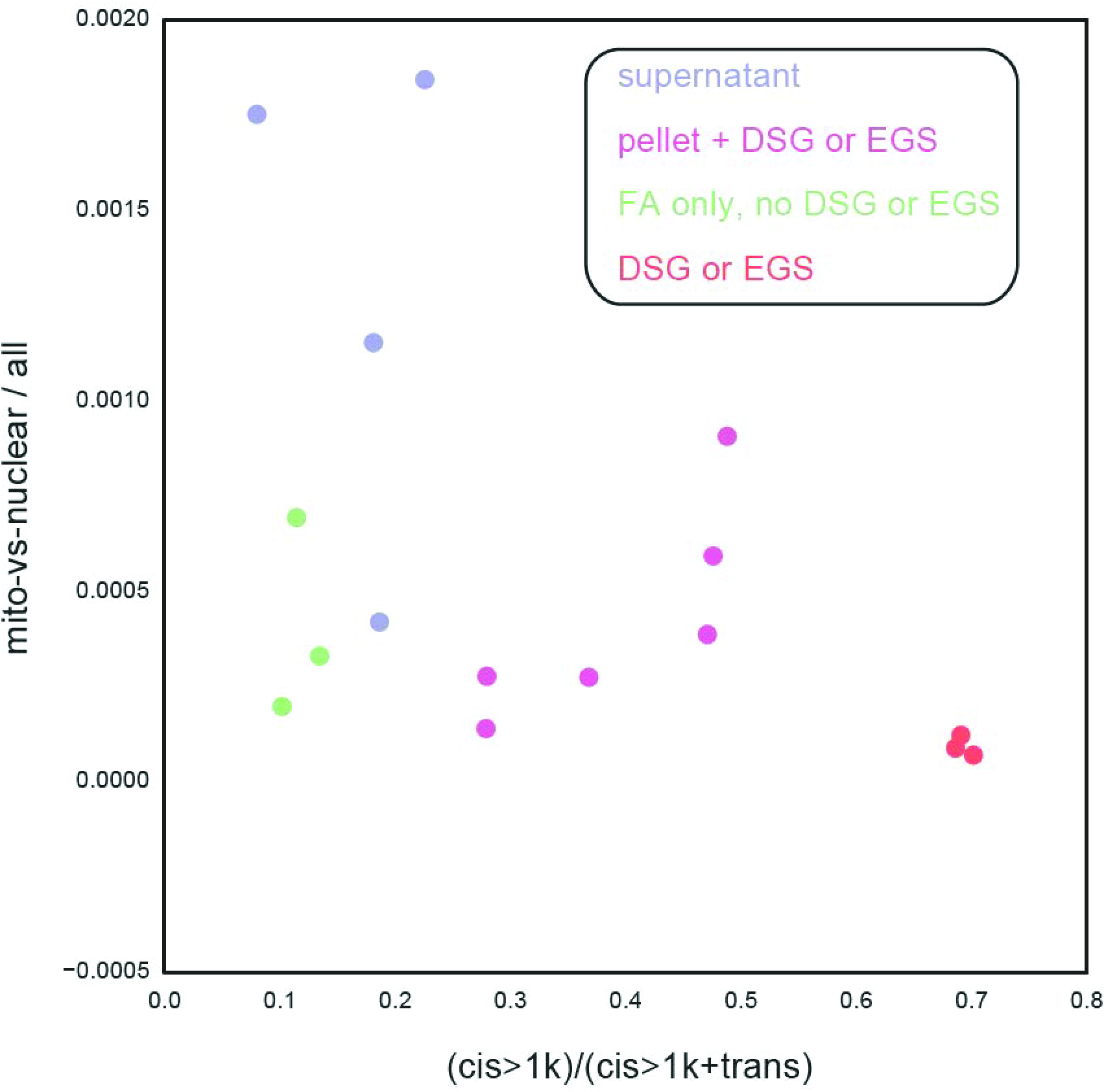
tRNA metagene analysis. Data for *S. cerevisiae* DSG pellet are shown here aligned for all tRNA genes in the yeast genome. (**a**) shows data normalized only for sequencing depth, while (**b**) shows data following matrix balancing. In each case left panel shows interaction counts, while right panel shows observed/expected relative to interaction distance.

**Fig. S16.**
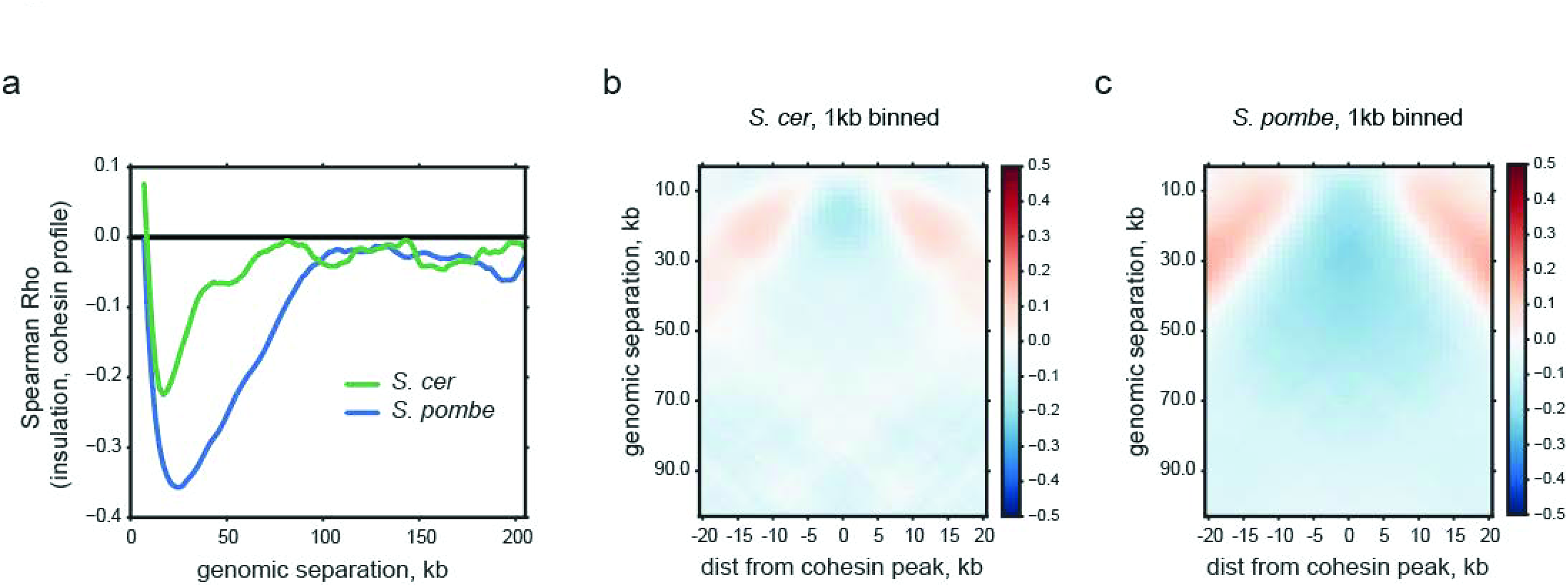
Cohesin insulates chromatin domains from one another. (**a**) Correlation between cohesin localization and local chromatin insulation (y axis), at varying offsets (x axis). Here, insulation was calculated as using sliding diamond window, as in^46^. Insulation profiles were calculated from 1kb binned and corrected DSG pellet contact maps, using a 10kb sliding window at each indicated offset. Note that cohesin localization is correlated with the local insulation score in both budding and fission yeast, but that cohesin-associated insulation in fission yeast is far stronger (deeper peak), and extends over greater genomic distances (peak width). Cohesin localization for *S. Cerevisiae* was obtained from GEO GSE42655 (Scc1,^47^), divided by input, log2-transformed, and binned to the same 1kb resolution as contact maps. Cohesin for *S. Pombe was* obtained from GEO GSE56848 (Psc3 WT,^24^), log2-normalized by input, and binned to 1kb. (**b-c**) Metagene insulation profiles for Micro-C contacts surrounding cohesin binding sites in the indicated species. Although both species exhibit local insulation, seen here as a blue depletion of contacts centered on cohesin binding sites, the inhibition of crossing interactions occurs at far greater distances (up to ∼75 kb) in *S. pombe* than in *S. cerevisiae* (∼25 kb). Cohesin peaks were called as local maxima on the binned 1kb log2 profile, and were additionally required to have a minimum spacing of 10kb and be in the top 75^th^ percentile overall. While displaying qualitative similarities, the quantitative differences captured in (**a-c**) may point to important differences in the underlying biology of cohesin in these two highly diverged yeast species. In particular, cohesin has been reported not to display peaks along the chromosomal arms in S. *cerevisiae* G1^48^, whereas peaks of cohesion binding in *S. pombe* G1 have been reported to coincide with regions of local insulation in *S. pombe* G1 Hi-C maps^24^. Together, these observations point towards a more important role for cohesin in organizing the arms of *S. pombe* chromosomes in interphase, relative to *S. cerevisiae.*

**Fig. S17.**
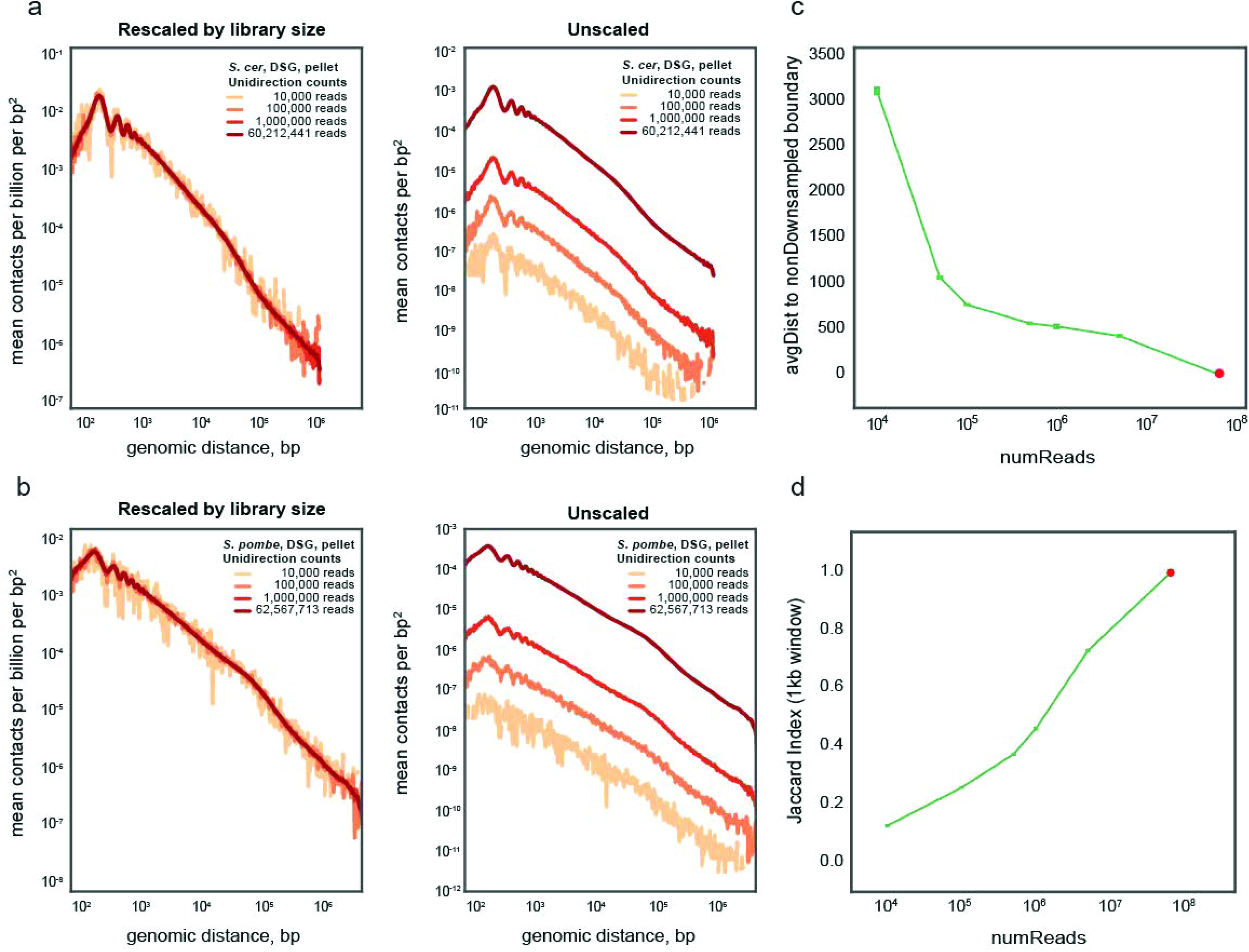
Subsampling Micro-C data. (**a-b**) Plots of interaction vs. distance are extremely robust to downsampling of sequencing data. Left plots show normalized density of interactions per squared basepair (y axis, normalized to total number of reads) vs. distance (x axis) for data downsampled to 10,000, 100,000, or 1,000,000 reads, or for the entire dataset. Right panels show the four curves separately, without normalization to sequence depth. In all cases, Micro-C XL reads (DSG pellet) were downsampled (after removing PCR duplicates) to the indicated number of reads. (**c**) Micro-C XL reads (*S. cerevisiae*, DSG pellet) were downsampled as indicated (x axis), and boundaries between CIDs were called as previously described^17^. Y axis shows distance between the boundary location called in the downsampled dataset and the nearest boundary called from the full dataset. Curves show average over ten subsampling analyses, and squares at each value of reads indicate the small standard deviation of these replicas. Note that even with only 100,000 sequencing reads, chromatin boundaries are identified to within 1 kb. The red circle represents the full dataset, where the average distance is zero. (**d**) Downsampling results in monotonic loss of boundary information. Here, y axis shows the average Jacard index of CID boundaries called from the full dataset versus the downsampled dataset with the number of reads indicated on the x axis. Curves again show the average of ten subsampling analyses, and the red circle represents the full dataset, where the Jaccard index is one. We note that the performance of this particular boundary-calling method does not represent a fundamental limit on the recovery of domain boundaries from sparse datasets, which represents a possible topic for future computational methods.

**Table S1. Read numbers and interactions vs. distance for all datasets.** Each sheet includes data for varying types of interactions (convergently oriented “IN-IN” read pairs, etc.) at varying distances (Column B) for all Micro-C datasets in this manuscript.

## Micro-C XL Protocol

### I. First Crosslinking

1. Culture 100 mL of yeast to the midlog stage, OD=0.55 o/n.
2. Add 37% formaldehyde directly to the culture to 3% of a final concentration.
3. Shake the culture at 210 rpm for 15 min at 30°C (<u>FA only</u>) or 10 min at 30°C (<u>Dual crosslinking</u>).
4. Quench the crosslinking by adding 10 mL of 2.5M Glycine.
5. Incubate for 5 min at room temperature.
6. Centrifuge the cells at 4000 rpm for 5 min at 4°C.
7. Pour off the medium and wash the cells in 50 mL of sterile water by vortexing.
8. Centrifuge the cells at 4000 rpm for 5 min at 4°C.
9. Pour off the water.

### II. Permeabilize the cell wall

1. Resuspend the cell pellet in 10 mL of Buffer Z and add 7 µL of 2-Mercaptoethanol (final 10mM).
2. Add 250 µL Zymolyase solution (final 250 µg/mL).
3. Shake the tube at 210 rpm for 40 min at 30°C.
4. Centrifuge the cells at 4000 rpm for 10 min at 4°C.
5. Aspirate the supernatant with a vacuum suction.
6. Rinse the permeabilized cells by 5mL cold 1X PBS.
7. Centrifuge the cells at 4000 rpm for 2 min at 4°C.
8. Aspirate the supernatant with a vacuum suction.

### III. Second Crosslinking

1. Freshly prepare the long crosslinker stock and working solution as below:

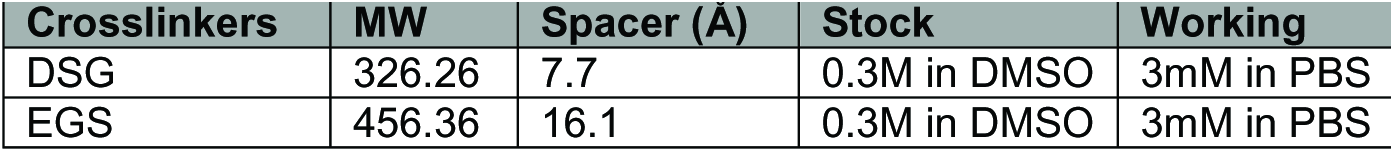
2. Resuspend the cells homogenously by 5 mL of working solution.
3. Rotate the tube for 40 min at 30°C.
4. Quench the crosslinking by adding 1 mL of 2.5M Glycine.
5. Centrifuge the cells at 4000 rpm for 10 min at 4°C.
6. Aspirate the supernatant with a vacuum suction.
7. Rinse the permeabilized cells by 5 mL cold 1X PBS.
8. Centrifuge the cells at 4000 rpm for 2 min at 4°C.
9. Aspirate the supernatant with a vacuum suction.
10. The crosslinked pellet can be store at -80°C for few months.

### IV. Chromatin fragmentation

1. Resuspend the cell pellet in 200 µL of MBuffer#1 (<u>freshly complete</u>).
2. Add the appropriate amount of MNase to digest the chromatin to > 95% mononucleosomes.
3. Incubate the tube for 20 min at 37°C.
4. Add 2 mM EGTA and incubate the tube for 10 min at 65°C to stop the MNase activity.
5. Here, you can proceed the sample depending on the experimental design: 1) Total chromatin. 2) Supernatant. 3) Pellet. Any of those fractions can be subjected to following Micro-C protocol.

### V. Chromatin cleaning and concentration

#### 1) Total chromatin

1. Transfer the whole sample into the 0.5 mL Amicon 10K spin column.
2. Concentrate the sample at 16000xg for at 4°C until the volume goes down to ∼50 µL.
3. Wash / pipette the sample by 450 µL MBuffer#2.
4. Repeat wash step 2 - 3.
5. Concentrate the sample at 16000xg for at 4°C until the volume goes down to ∼30 µL.
6. Add BSA to 1X final concentration.

#### 2) Supernatant

1. Centrifuge the tube at 16000xg for 5 min at 4°C.
2. Collect the supernatant.
3. Concentrate the sample by the 0.5 mL Amicon 10K spin column at 16000xg for at 4°C until the volume goes down to ∼50 µL.
4. Wash / pipette the sample by 450 µL MBuffer#2.
5. Repeat wash step 2 - 3.
6. Concentrate the sample at 16000xg for at 4°C until the volume goes down to ∼30 µL.
7. Add BSA to 1X final concentration.

#### 3) Pellet

1. Centrifuge the tube at 16000xg for 5 min at 4°C.
2. Collect the pellet.
3. Resuspend the pellet in 1 mL MBuffer#2.
4. Centrifuge the tube at 16000xg for 5 min at 4°C.
5. Aspirate the buffer with a vacuum suction.
6. Repeat wash steps 3 - 5.
7. Resuspend the pellet to 30 µL of MBuffer#2 + final 1X BSA (or NEBuffer 2.1).

### VI. Repair and label the end of chromatin fragments

#### 1) De-phosphorylation

| Total | 32.5µL | Final condition |
| --- | --- | --- |
| Chromatin sample | 30µL | 50mM NaCl, 10mM Tris, 10mM MgCl<sub>2</sub>, 1X BSA |
| 1U/µL r-Shrimp alkaline phosphatase | 2.5 | 2.5U |

→ Incubate for 45min at 37°C.
→ Inactivate for 5min at 65°C.

#### 2) End-Chewing

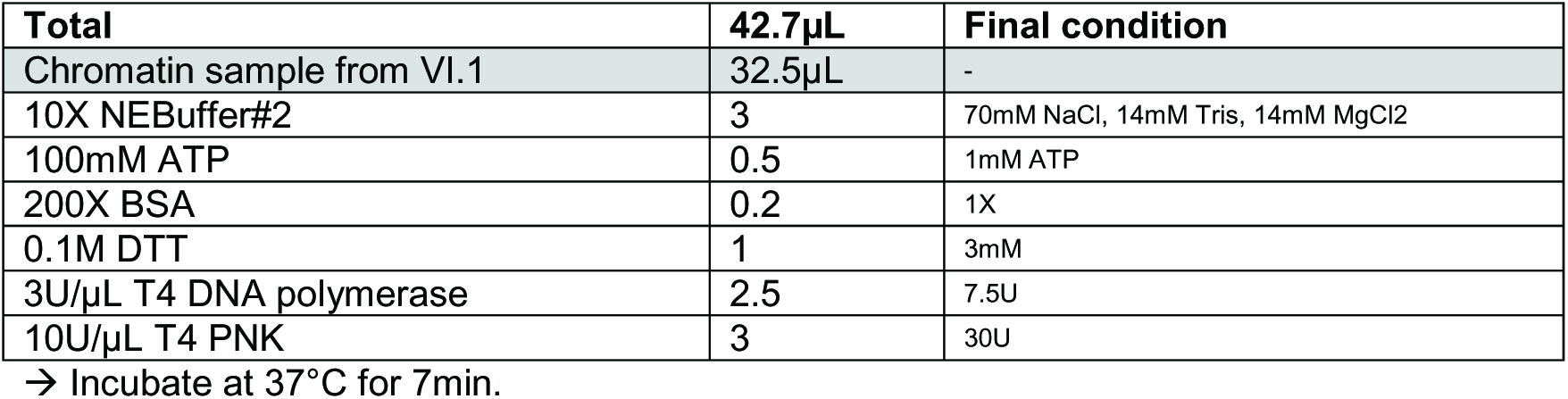

| Total | 42.7µL | Final condition |
| --- | --- | --- |
| Chromatin sample from VI.1 | 32.5µL | - |
| 10X NEBuffer#2 | 3 | 70mM NaCl, 14mM Tris, 14mM MgCl<sub>2</sub> |
| 100mM ATP | 0.5 | 1mM ATP |
| 200X BSA | 0.2 | 1X |
| 0.1M DTT | 1 | 3mM |
| 3U/µL T4 DNA polymerase | 2.5 | 7.5U |
| 10U/µL T4 PNK | 3 | 30U |

→ Incubate at 37°C for 7min.

#### 3) End-labeling

| Total | 100µL | Final condition |
| --- | --- | --- |
| Chromatin sample from VI.2 | 42.7µL | - |
| 0.4mM Biotin-dATP | 25 | 100µM |
| 0.4mM Biotin-dCTP | 25 | 100µM |
| 10mM dTTP + dGTP | 1 | 100µM |
| 10X T4 DNA ligase buffer | 6 | 30mM NaCl, 35mM Tris, 12mM MgCl<sub>2</sub>, 7mM DTT |
| 200X BSA | 0.3 | 1X BSA |

→ PCR machine: Incubate for 25min at 25°C → 15min at 12°C → 4°C.

→ Add EDTA (final 30mM) and heat inactivation for 20 min at 65°C.

### VII. Proximity ligation and Remove unligated ends

#### 1) Ligation

*Although in our test Micro-C in “pellet” can be scaled down to 1mL ligate reaction, we suggest at least using 2.5 mL for routine experiment.*

| Total | 2.5mL<br>(pellet) | 10mL<br>(Total & Sup) | Final condition |
| --- | --- | --- | --- |
| Chromatin sample from VI. | 100uL | 100μL | - |
| Water | 2122 | 8809 | - |
| 10X T4 DNA ligase buffer *w/ ATP* | 250 | 1000 | 1X |
| 200X BSA | 12.5 | 50 | 1X |
| 1M MgCl<sub>2</sub> | 3 | 3 | Equal to the moles of EDTA from previous part. |
| 400U/μL T4DNA ligase | 12.5 | 38 | 1.5 – 2U/μL |

- Incubate for 60min at room temperature. - **Pellet:** Centrifuge the pellet by 16000xg for 10min at 4°C. - **Total/Sup:** 15mL Amicon 30k concentrates sample by spin at 4000g for 40min at 4°C.

#### 2) Remove the biotin-dNTP at unligated ends

| Total | 100μL<br>(pellet) | 280μL<br>(Total & Sup) | Final |
| --- | --- | --- | --- |
| Chromatin sample | - | 250 | - |
| 10X NEBuffer#1 | 10 | 28 | 1X |
| Water | 89 | - | - |
| 100U/μL Exonuclease III | 1 | 1 | 100U |

- Incubate for 5 min at 37°C.

#### 3) Reverse crosslinking

→ Add 20X proteinase K to 1X final concentration.

→ Incubate for overnight at 55°C.

### VIII. Dinucleosomal DNA purification

1. Phenol:Chloroform:Isoamyl Alcohol extraction twice → spin at 19800xg for 10min.
2. Ethanol precipitation: 0.1x volume of sodium acetate and 2.5X volume of 100% ethanol → -80°C for > 1hr → spin at 19800xg for 15min at 4°C → wash pellet by 75% ethanol → spin at 19800xg for 5 min at 4°C → Air dry pellet for 10min.
3. Resuspend pellet in 50uL of TE buffer (+ 1X RNase solution) and incubate for 30min at 37°C.
4. ZymoClean to purify DNA.
5. Run DNA samples on 3% Nusieve agarose DNA gel.
6. Size selection of the band between 250 – 350 bp.
7. ZymoGel purification and dissolve final product in 17 µL of elution buffer.
8. Quantify the input DNA by Qubit.

### IX. Library construction by “with-bead” method

#### 1) End-it

| Total | 25μL |
| --- | --- |
| DNA | 17 |
| 10X End-it buffer | 2.5 |
| 10X ATP | 2.5 |
| 10X dNTP | 2.5 |
| End-it enzyme mix | 0.5 |

- Incubate for 45 min at room temperature.
→ 2X Ampure XP purification.

#### 2) A-tailing

|  |  |
| --- | --- |
| Total | 25µL |
| DNA | 16 |
| 10X Exo- Klenow buffer | 2.5 |
| 1mM dATP | 5 |
| Exo- Klenow fragment enzyme | 1.5 |

→ Incubate for 30min at 37°C.
→ 2X Ampure XP purification. (PEG/NaCl solution: 20% PEG, 2.5M NaCl)

#### 3) Adapter ligation

|  |  |
| --- | --- |
| Total | 15µL |
| In-line / Indexing Adapter | Ratio of Adapter:Input DNA = 10:1 – 50:1 |
| Water | to total 15µL |
| 10X Fast-link DNA ligase buffer | 1.5 |
| 10X ATP | 1.5 |
| Fast-link DNA ligase | 1 |

→ Incubate for > 2 hr at room temperature.
→ Add 10µL of EB to total volume 25 µL.
→ 2X Ampure XP purification and elute DNA in 150 µL of water.

#### 4) Streptavidin beads purification

1. Wash 2.5 µL of beads per sample (100mL culture) by 1X TBW twice.
2. Resuspend the washed beads in 150 µL 2X BW.
3. Mix with 150 µL of adapter-ligated DNA sample.
4. Rotate for 15 min at room temperature.
5. Wash by 500 µL of 1X TBW twice.
6. Rinse by 200 µL of MBuffer#2.
7. Resuspend in 15 – 20 µL of EB buffer.

#### 5) On-beads PCR

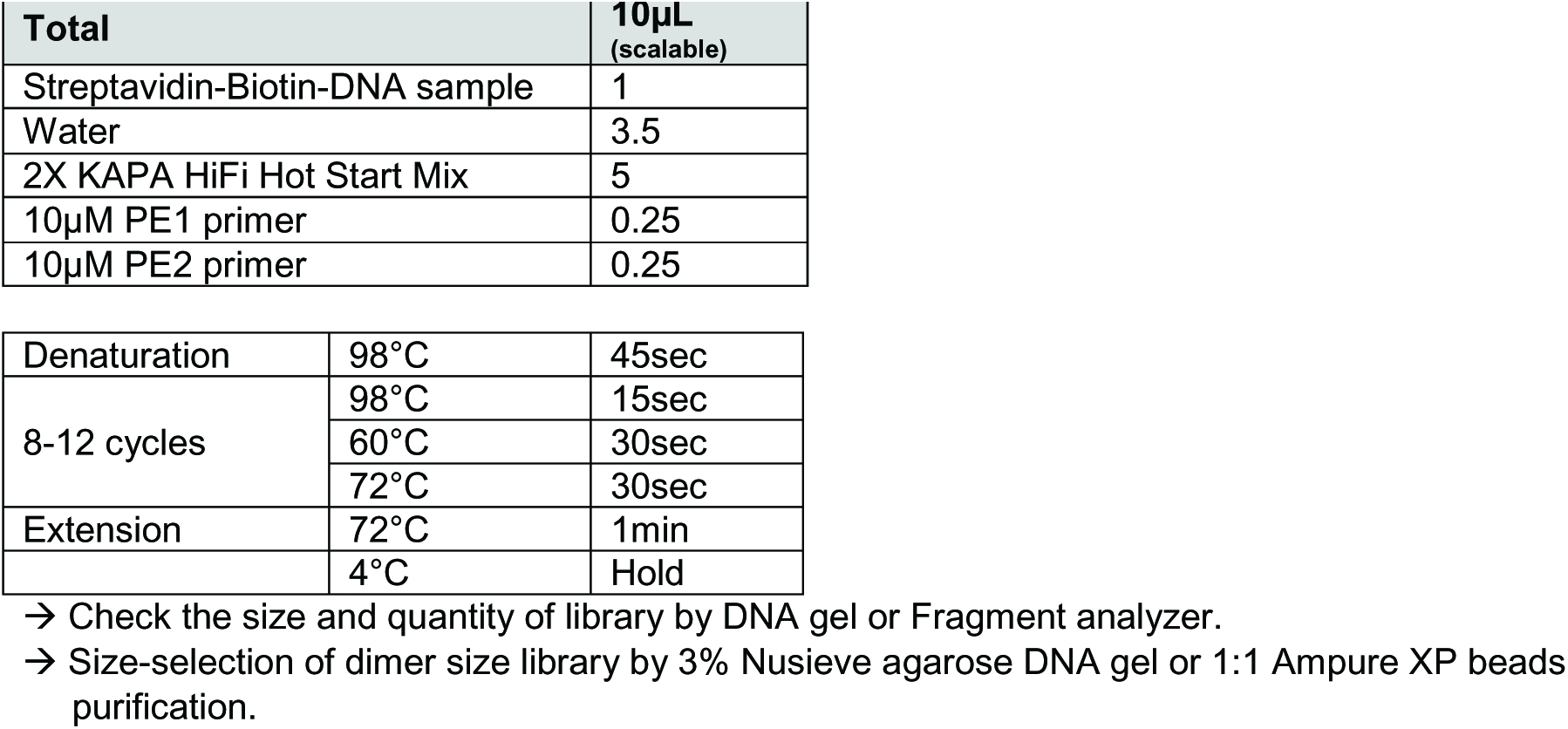

| Total | 10µL<br>(scalable) |
| --- | --- |
| Streptavidin-Biotin-DNA sample | 1 |
| Water | 3.5 |
| 2X KAPA HiFi Hot Start Mix | 5 |
| 10µM PE1 primer | 0.25 |
| 10µM PE2 primer | 0.25 |

|  |  |  |
| --- | --- | --- |
| Denaturation | 98°C | 45sec |
| 8-12 cycles | 98°C | 15sec |
|  | 60°C | 30sec |
|  | 72°C | 30sec |
| Extension | 72°C | 1min |
|  | 4°C | Hold |

→ Check the size and quantity of library by DNA gel or Fragment analyzer.
→ Size-selection of dimer size library by 3% Nusieve agarose DNA gel or 1:1 Ampure XP beads purification.

### X. Deep sequencing by Illumina PE-50

#### Solutions and Enzymes

- **YPD:** yeast extract/peptone/dextrose
- **37% Formaldehyde** (Sigma Aldrich # 252549)
- **DSG (disuccinimidyl glutarate)** (ThermoFisher #20593)
- **EGS (ethylene glycol bis(succinimidyl succinate))** (ThermoFisher #21565)
- **2.5M Glycine** (Sigma Aldrich #G7126)
- **Buffer Z:** 1M sorbitol, 50mM Tris pH 7.4
- **14.3M 2-Mercaptoethanol** (Sigma Aldrich # M6250)
- **Zymolyase solution:** 10mg/ml in Buffer Z; lasts up to 2 weeks at 4°C (Sunrise Science #N0766555)
- **MBuffer#1:** 50mM NaCl, 10mM Tris-HCl pH 7.5, 5mM MgCl_2_, 1mM CaCl_2_, and **freshly** complete with 0.5mM spermidine, 1mM B-ME, and NP-40 (percentage is determined by titrating the ratio of pellet / sup).
- **Micrococcal Nuclease** (Worthington Biochem): resuspended from lyophilized powder at 20 U/μl in Tris pH 7.4. Aliquot into tubes upon first use and freeze at –80°C.
- **0.5M EDTA** (Life technology # AM9261)
- **MBuffer#2 (NEBuffer#2):** 50mM NaCl, 10mM Tris-HCl pH 7.5, 10mM MgCl2
- **Shrimp Alkaline Phosphatase** (**rSAP**) (New England Biolabs # M0371)
- **T4 DNA Polymerase** (New England Biolabs # M0203)
- **T4 Polynucleotide Kinase** (New England Biolabs # M0201)
- **Biotin-14-dCTP** (Life Technologies # 19518018)
- **Biotin-14-dATP** (Life Technologies # 19524016)
- **T4 DNA Ligase** (New England Biolabs # M0202)
- **Exonuclease III (*E. coli*)** (New England Biolabs # M0206)
- **20X Proteinase K solution:** TE with 20 mg/ml proteinase K and 50% glycerol. Store in -20°C.
- **Elution buffer (EB):** 10 mM Tris-HCl pH 7.5
- **TE buffer:** 10 mM Tris-HCl pH 8.0, 1 mM EDTA
- **End-It DNA End-Repair Kit** (EpiCentre BioTechnologies # ER81050)
- **Exo-Minus Klenow DNA Polymerase** (EpiCentre BioTechnologies # KL111)
- **Fast Link DNA Ligation Kit** (EpiCentre BioTechnologies # lk6201)
- **Dynabeads^®^ MyOne Streptavidin C1** (Life Technologies # 65001)
- **KAPA HiFi HotStart ReadyMix** (KAPA Biosystems # KK2601)

